# Structural variation affecting DNA backbone interactions underlies adaptation of B3 DNA binding domains to constraints imposed by protein architecture

**DOI:** 10.1101/2020.06.05.137281

**Authors:** Haiyan Jia, Masaharu Suzuki, Donald R. McCarty

**Affiliations:** Horticultural Sciences Department, Plant Molecular and Cellular Biology Program, University of Florida, Gainesville, Florida, 32611-0690, USA

## Abstract

Functional diversification of transcription factor families through variation in modular domain architectures has played a central role in the independent evolution of gene regulatory networks underlying complex development in plants and animals. Here we show that architecture has in turn constrained evolution of B3 DNA binding domains in the B3 network regulating embryo formation in plants. B3 domains of ABI3, FUS3, LEC2 and VAL1 proteins recognize the same cis-element. ABI3 and VAL1 have complex architectures that physically integrate cis-element recognition with other signals, whereas LEC2 and FUS3 have reduced architectures conducive to their roles as pioneer activators. Qualitatively different activities of LEC2 and ABI3 B3 domains measured in vivo and in vitro are attributed in part to clade-specific substitutions in three amino acids that interact with the DNA backbone. Activities of FUS3 and VAL1 B3 domains show a similar correlation with architectural complexity. Domain-swap analyses in planta show that in a complex architecture setting the attenuated activities of ABI3 and VAL1 B3 domains are required for proper integration of cis-element recognition with hormone signalling. These results highlight modes of structural variation affecting non-specific, electrostatic interactions with the DNA backbone as a general mechanism allowing adaptation of DNA binding affinity to architectural constraints while preserving DNA sequence specificity.

## INTRODUCTION

In plant and animal lineages that have independently evolved embryos (1), key gene regulatory networks that underlie complex development have evolved through functional diversification within families of related transcription factors (2). Functional diversification is facilitated by the typically modular domain architectures of transcriptional regulators. In plants, transcription factors that form a network controlling embryo formation and subsequent transition to the vegetative development share a conserved B3 DNA binding domain (3).

In the B3 network of Arabidopsis, the AFL (ABI3, FUS3 and LEC2) B3 domain proteins primarily function as activators that promote embryogenesis (3), whereas VAL1-type B3 proteins repress the AFL network prior to germination of the seed enabling a transition to vegetative development (4,5). While the B3 domains of AFL activators (3) and VAL repressors bind specifically to the same Sph/RY cis-element (6–11), their distinct roles in the network are differentiated in part at the level of protein domain architecture. ABI3 and VAL1 have complex, multiple domain architectures that physically integrate B3 recognition of the Sph/RY cis-element with hormonal signals (3)and chromatin marks (3,6,7,11–13), respectively, whereas LEC2 and FUS3 have architectures of reduced complexity conducive to their roles as pioneer activators (14,15). Orthologs of VP1/ABI3 and VAL1 type proteins occur in all sequenced land plant genomes (16), whereas FUS3 and LEC2 genes are restricted to seed plants and rosid clade of angiosperms, respectively (16). This phylogenetic relationship is consistent with FUS3 and LEC2 proteins having evolved via truncation of VP1/ABI3 type ancestors (15,16). While the DNA sequence specificities of B3 domains in VAL1 and AFL proteins are conserved, the contributions of intrinsic differences in B3 domain properties to functional diversification in the B3 transcription factor network have not been systematically addressed.

Here we show that intrinsic differences in activities of ABI3 and LEC2 B3 domains, measured both in yeast and in a transgenic plant assay that captures integration with abscisic acid (ABA) signaling in a complex architecture setting, reflect a pattern of B3 domain adaptation to complex and reduced protein architectures that extends to VAL1 and FUS3. We identify a clade-specific triad of amino acid substitutions implicated in B3 domain interaction with the DNA backbone as a structural basis for functional differentiation of LEC2 and ABI3 B3 domains. We suggest that modes of structural variation that affect non-specific DNA backbone interactions while preserving sequence specific base contacts provide a general mechanism for adapting a conserved DNA binding domain to diverse architectural environments that arise in the evolution of gene regulatory networks.

## MATERIALS AND METHODS

### Structural analysis and modeling

Structural models of the ABI3 B3 domain bound with DNA were made by MODELLER v9.20 (17) using crystal structures of VAL1 (PDB ID: 6J9A, 6FAS) and FUS3 B3 (PDB ID: 6J9B) complexed to Sph/RY DNA as templates. Fifty replicate structural models were constructed for each template. The best model was selected based on the objective function score (17). Structure analyses and molecular graphics were done in UCSF Chimera (18) (https://www.cgl.ucsf.edu/chimera).

### Phylogenetic analysis

Phylogenetic analyses were performed using online tools hosted at NGPhylogeny (https://ngphylogeny.fr/). Multiple protein sequence alignments of B3 domains were made using MAFFT (19)and maximum-likelihood phylogenetic trees were generated using PHYML (20). Bootstrap values were calculated using BOOSTER (21).

### Yeast One hybrid (Y1H) assays

To produce the WT AFL and VAL Y1H effector plasmids, the B3 domains of ABI3, FUS3, LEC2, VAL1, VAL2 and VAL3 were amplified by PCR (PrimeSTAR TM HS DNA Polymerase, TaKaRa) using the plasmids carrying the full-length cDNA of each gene as template. The corresponding PCR products were cloned into the pGADT7 vector (Clontech, CA) between the *ClaI* and *SacI* sites, which generated N terminal GAL4 activation domain (GAL4 AD) fusion for each Y1H effector. pGADT7 has the *LEU2* selection marker in yeast.

The B3 domain mutant effector plasmids were produced by using AFL WT B3 domain effector plasmids as templates and then introducing point mutations at the 3 targeted residues (residues 64,66,and 69), respectively. All site-directed mutagenesis reactions were performed by following the instructions in the QuikChange Lightning Multi Site-Directed Mutagenesis Kit (Agilent Technologies, Stratagene, CA). Primers used for PCR and site-directed mutagenesis reactions are indicated in Supplemental Table S2. The Y1H reporter plasmid *pHISi-1-Sph2* was constructed by inserting a 40bp fragment that contains Sph dimer (named as Sph2) into the pHISi-1 vector (with *HIS3* selection marker): 5’-GAT<u>CATGCAT</u>GGACGACACGGAT<u>CATGCAT</u>GGACGACACG-3’ (Sph underlined). The pHISi-1 vector was provided by the Matchmaker one-hybrid kit (Clontech, CA). The 40 bp promoter sequence was originally cloned from the maize *C1* gene promoter (22). All the constructs were verified by sequencing.

To generate the reporter strain Sph2-YM4271, Y1H reporter plasmids pHISi-1-Sph2 were linearized by *XhoI* and stably integrated into the non-functional *HIS3* locus of Yeast YM4271 by homologous recombination following the manufacturer’s instructions (Clontech, CA). To control basal expression of *HIS3* reporter gene, Sph2-YM4271 cells were grown on synthetic dropout (SD)-His media supplemented with the inhibitor 3 mM 3-amino-1,2,4-triazole (3-AT; Sigma). All Y1H effector plasmids were separately transformed into the Sph2-YM4271 reporter strain and plated on SD/-Leu plates. The plates were incubated at 30°C for 4 days to screen the positive transformations.

To test the interaction of effectors with the sph2 reporter, transformed yeast reporter cells were cultured overnight in SD-Leu liquid medium. Cells were then pelleted, washed and resuspended in water. OD_600_ was measured using a microplate reader Epoch (Biotek). Then the concentration of each transformed yeast cell culture was normalized to OD_600_ = 0.04. Ten-fold serial dilutions of each normalized culture were prepared. A 3 μl sample from each culture in the dilution series was spotted on SD/-Leu, SD/-Leu/-His, and SD/-Leu/-His plus 3mM 3-AT plates, respectively, and incubated at 30°C for 4 days.

For yeast growth curves, cell cultures that normalized to a uniform OD_600_ (OD_600_ = 0.04) in 1.5ml of SD/-Leu/-His liquid medium (plus 3 mM 3-AT) were incubated in a 12-well microplate (Thermo Scientific) at 30°C for 96 hours (h). The OD_600_ was measured in a microplate reader Epoch (Biotek) at 12 h intervals. 3 replicates of each treatment were used in these experiments.

### Construction of *VP1::B3* chimeral transgenic constructs

The *35S-VP1* construct was created previously in our lab (23). The full length *VP1* cDNA that includes *XbaI* and *XhoI* sites at the 5’- and 3’-ends, respectively, was amplified using the *VP1-XbaI* / *VP1-XhoI* primer pair (Supplemental Table S3) and *35S-VP1* plasmid as a template. The resulting PCR product was sub-cloned into pCR4-TOPO vector (Invitrogen) to create VP1-TOPO. AFL B3 and VAL1 chimeral transgene *(Pro35S:VP1::ABI3-B3, Pro35S:VP1::FUS3-B3, Pro35S:VP1::LEC2-B3 and Pro35S:VP1::VAL1-B3)* constructs were produced by the following several steps: 1) *EcoRV* and *SacI* restriction sites that flanked the B3 domain were introduced into the *VP1-TOPO* plasmid by site-directed mutagenesis (Agilent Technologies, CA); 2) The plasmids created in step 1 and the respective PCR amplified AFL and VAL1 B3 domain sequences that carrying *EcoRV* and *SacI* at both ends were digested with *EcoRV*and *SacI*. The restriction fragments containing the VP1-TOPO plasmid backbone and the AFL and VAL1 B3 domain sequence were purified and ligated together; 3) The restriction sites at the swap borders were then restored to the original sequence by site-directed mutagenesis; and 4) AFL B3 and VAL1 chimeral transgenes *(VP1::ABI3-B3, VP1::FUS3-B3, VP1::LEC2-B3* and *VP1::VAL1-B3)* in TOPO plasmid were digested with *XbaI/XhoI* and ligated into the *XbaI/SalI* sites of the transformation vector *pCAMBIA1300*, which placed the 35S promoter upstream of the transgene and attached a GFP tag to C-terminus of the hybrid protein (Figure 2A). The mutant chimeral transgenes were created by site-directed mutagenesis using corresponding templates in the TOPO plasmids and ligated to *pCAMBIA1300*. The Primers used for PCR and site-directed mutagenesis reactions are indicated in Supplemental Table S3. All the constructs were verified by sequencing.

### Generation of transgenic plants

The wild type and mutant chimeral transgenic constructs were used to transform to agrobacterium strain *GV3101*, and plated on YEP media (10g yeast extract, 10g Bacto peptone, and 5g NaCl, pH =7.0) containing 50mg/L Kanamycin (Kan) and 25mg/L Gentamicin sulfate. Agrobacterium-mediated transformation of *abi3-6* mutant Arabidopsis plants was performed by using the floral-dip method (24). T1 seeds were harvested about 3-4 weeks after transformation. T1 seeds were sterilized and stratified at 4°C in dark for 3 days. To screen for hygromycin resistance, stratified T1 seeds were plated on MS media containing 1X Murashige and Skoog salt, 0.05% MES, 1% sucrose sterilized by filtration, and 0.15% of phytagel (Sigma) supplemented with 25mg/L hygromycin and 200mg/L Carbenicillin (to kill the Agrobacterium), and grown at 23°C for one week under continuous light. Plants that exhibited rapid relative growth rates were selected as candidate transgenic seedlings and grown on the MS media containing 25mg/L Hygromycin for an additional week. The surviving hygromycin resistant T1 seedlings that had good root formation were transferred to soil using plant growth conditions described previously (5). T2 seeds of independent lines that segregated approximately 3:1 brown: green colored seeds were screened again for hygromycin resistance. The homozygous transgenic seeds were screened in the T3 generation by scoring seed color (all brown) as well as 100% Hygromycin resistance on selection medium.

### Gene expression analysis

Detached leaf tissue of 14-d-old T1 transgenic seedlings were incubated 24h on MS and MS plus 5 μM ABA, respectively. Col-0 WT and *abi3-6* seedlings were used as controls in this assay. Total RNA was extracted from the leaf samples, or whole T1 seedlings in cases of abnormal morphology, using the plant miniRNA kit (Zymo research). All the conditions for quantitative real-time RT-PCR and RT-PCR were as described previously (5). *Tub2 (AT5G62690)* was used as an endogenous control in the RT-PCR. Transgene expression was quantified using primers that targeted the NOS terminator region common to all transgenes. The primers used for RT-PCR and quantitative PCR are listed in Supplemental Table S3.

### *In vitro* DNA binding assays

The AFL and VAL1 B3 domains were sub-cloned by using *VP1::B3* chimeral transgenic constructs (Supplemental Figure S9) as templates. A common pairs of primers *B3-BamHI*/*B3-EcoRI* (Supplemental Table S4) were used to incorporate *BamHI* and *EcoRI* sites for cloning into pGEX-2T (pPhamacia Inc., Uppsala, Sweden). Constructs were verified by sequencing. All GST-B3 fusion proteins were expressed in *E.coli* BL21(DE3) cells and were purified by gluthathione-Sepharose affinity chromatography according to the manufacturer’s recommendations (GE Healthcare). The proteins were eluted in elution buffer (20 mM reduced glutathione, 100 mM Tris-HCl, pH 8.0). The purified GST fusion proteins were visualized by sodium dodecyl sulphate-polyacrylamide gel electrophoresis (SDS-PAGE). Protein concentrations were measured via bradford assay (25). Sph2 probe is 5’ biotin labelled (synthesized by IDT) and is shown in Supplemental Table S4.

B3-DNA binding was measured in a label-free in vitro kinetics assay at pH 7.0 using Octet Qke system (Pall ForteBio) (26). The 1X Kinetics Buffer (Pall ForteBio) was used as running buffer. The experimental steps are: after a 5 min initial baseline step, 0.05 μM biotinylated Sph2-probe was loaded onto streptavidin biosensors for 5 min until reach saturation. Probes were then quenched by 25 μg/ml biocytin for 2 min. After a 2 min baseline step, each probe was then exposed to B3 protein at increasing concentrations for 15 min at association step, followed by 30 min dissociation step. Changes in the number of molecules bound to the biosensor causes a shift in the interference pattern that is measured in real time (Pall ForteBio application note 14). Measurements were taken from one experimental replicate. The association rate (k_on_), dissociation rate (k_off_), kinetic K_d_ and steady state K_d_ were obtained by Octet Data Analysis HT software fitting a one-site binding curve using non-linear regression. Similar results were obtained from two experimental replicates.

In the gel mobility shift assay, DNA binding reactions contained 1fmol of Sph2-probe and various concentration of proteins suspended in a 15ul of binding reactions (LightShift Chemiluminescent EMSA kit, Catolog # 20148, Thermo Scientific). The components in the reactions [1X Binding buffer, 50ng/ul Poly (dI.dC), 2.5% of Glycerol, 5mM MgCl2, 0,05% NP-40]. The reactions were incubated at room temperature for 20 min and then resolved on 6% polyacrylamide gel run in 0.5 X TBE buffer at 4 °C. The remaining steps were performed as described in the LightShift Chemiluminescent EMSA kit (Thermo Scientific). The images were captured by a CCD camera or X-ray film. Images were analyzed by densitometry using image J.

### Chromatin immunoprecipitation-quantitative PCR (ChIP-qPCR) assays

ChIP assays were performed as the protocol described by Komar et al., (27). Briefly, 1.5 g Arabidopsis leaf tissue was fixed with 1% formaldehyde under vacuum and was grinded with liquid nitrogen. After nuclei were extracted and lysed, chromatin was fragmented to most abundant size at ~500 bp by Bioruptor 300 (Diagenode) for 30 cycles with the settings at high 30 sec ON/30 sec OFF at 4°C. The supernatant was pre-cleaned by incubation with Protein A/G MagBeads (Cat. No. L0027, Genescript) at 4 °C for 1 hour, and then immunoprecipitated by anti-GFP antibody (Cat. No, 50430-2-AP, Proteintech) coated protein A/G magnetic beads at 4 °C overnight. A mock sample without any antibody was prepared simultaneously. After reverse-crosslink, the precipitated DNA was purified and used as the templates for qRT-PCR with Luna Universal qPCR master mix (M3003G). The primer sequences were shown in Supplemental Table S5.

## RESULTS AND DISCUSSION

### Architectural diversification of B3 domain transcription factors

The VAL1 and VP1/ABI3 type B3-domain transcription factors with characteristic domain compositions occur in diverse vascular plant genomes indicating that these architectures arose in concert with evolution of land plants (embryophytes) consistent with their fundamental roles in plant embryo development. By contrast, FUS3- and LEC2-type transcription factors, which have similar architectures of reduced complexity, are restricted to seed plants and to the rosid clade of the angiosperms (15,16), respectively, suggesting that they evolved via truncation of VP1/ABI3-like ancestors.

#### Evidence for architectural constraints on B3 domain evolution

A phylogenetic analysis of B3 domain sequences was undertaken to further resolve relationships among the B3 domains of VAL1 and AFL proteins (Figure 1A and Supplemental Figure S1). A salient feature of the B3 domain tree is that the VP1/ABI3 clade, which spans ~400 MY of land plant evolution, is clearly separate from the putatively derived FUS3 and LEC2 B3 clades. This separation is noteworthy because if 1) FUS3 and LEC2 proteins were indeed derived from ancestors in the VP1/ABI3 clade, and 2) conservation of DNA binding sequence specificity were the sole functional constraint on B3 domain evolution, then we would instead expect FUS3 and LEC2 B3 clades to be nested within the VP1/ABI3 clade consistent with their later origins within the seed plants. To account for the observed separation of VP1/ABI3 and FUS3/LEC2 B3 clades, we hypothesized that evolution of B3 domains in VP1/ABI3 type proteins is subject to additional functional constraints imposed by a complex architecture. A key prediction of that hypothesis is that B3 domains derived from proteins with simple and complex architectures, respectively, are not functionally equivalent.

**Figure 1.**
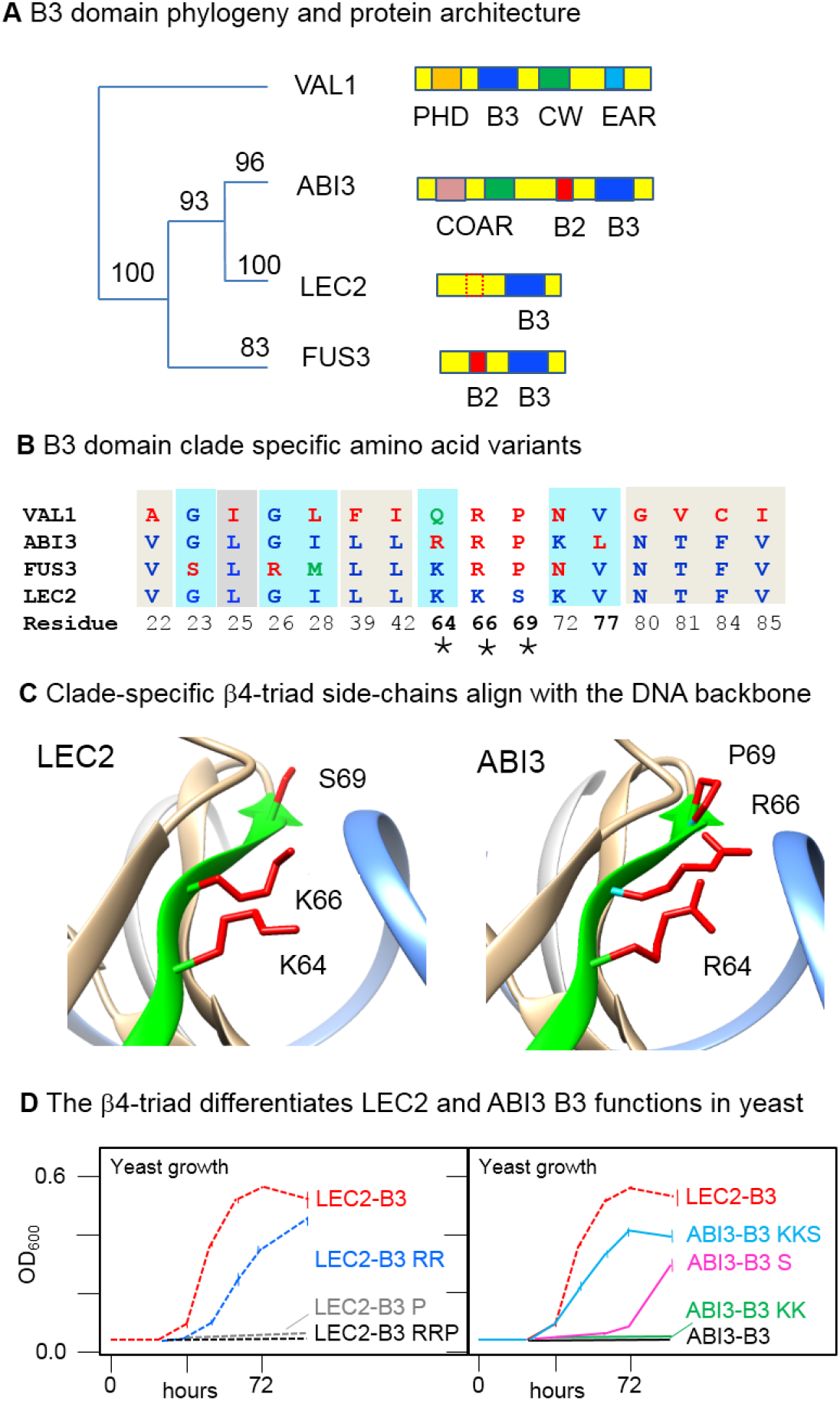
A structural basis for functional differentiation of LEC2 and ABI3 B3 domains. (**A**) B3 domain phylogeny and protein domain architectures of ABI3, FUS3, LEC2 and VAL1 transcription factors. The tree is adapted from a phylogenetic analysis of B3 domain sequences from diverse orthologs of ABI3, FUS3, LEC2 and VAL1 (Supplemental Figure S1). Multiple protein sequence alignments and maximum-likelihood tree were constructed using MAFFT (19) and PhyML (20) with bootstrap support based on BOOSTER (21). (**B**) B3 domains are distinguished by clade-specific structural variants at 16 amino acid positions. Clade-specific variant positions meet two criteria: 1) positions include an amino acid substitution in at least one B3 clade, and 2) all substitutions observed at that position are strictly conserved within each of the four clades. Here and elsewhere, amino acid positions are numbered uniformly according to the sequence alignment shown in Supplemental Figure S2. Eight clade-specific variants that distinguish VAL1 from the three AFL B3 clades are shaded in grey background. Six clade-specific variant positions that distinguish FUS3 B3 from ABI3 B3 are shaded light blue. Four clade-specific positions that distinguish ABI3 and LEC2 B3 sequences are highlighted with bold numbers. The latter include the β4-triad (marked by *), comprised of variants at positions 64,66, and 69 located in beta sheet strand 4 (Supplemental Figure S2). (**C**) Locations of β4-triad amino acids in the B3 domain align with the DNA backbone. Images showing β4-triad amino acid side-chains (colored red) in LEC2 B3 (6J9C.pdb) (9) and a structural model of ABI3 B3 based on FUS3 (6J9B.pdb) (9) were generated with Chimera (18). A ribbon depicting the β4 strand is colored green, and proximal DNA backbone strand is colored light blue. The ABI3 B3 model was constructed by MODELLER (17). (**D**) β4-triad substitutions differentiate *in vivo* activities of LEC2 B3 and ABI3 B3 domains in yeast. A yeast-one-hybrid (Y1H) assay for functional analysis of B3 domains is based on activation of a HIS selectable marker controlled by a minimal promoter containing a dimer of the Sph/RY cis-element recognized by AFL and VAL1 B3 domains (3,6–11) (Supplemental Figure S3). Activities of LEC2 B3 and ABI3 B3 effector proteins were quantified by measuring HIS dependent growth of yeast cultures (OD_600_, optical density at 600 nm) over a 96-hour time course. LEC2-B3: wildtype B3 fused to GAL4 activator (red, dashed); LEC2-B3 RR: K64R and K66R substitution mutations of the LEC2-B3 effector (blue, dashed); LEC2-B3 RRP: K64R, K66R, S69P triple substitution mutation of the LEC2-B3 effector (black, dashed); LEC2-B3 P: S69P substitution mutation of the LEC2-B3 effector (grey, dashed); ABI3-B3: wildtype B3 fused to GAL4 activator (black, solid); ABI3-B3 KK: R64K and R66K substitution mutation of the ABI3-B3 effector (green, solid): ABI3-B3 S: P69S substitution mutation of the ABI3-B3 effector (red, solid); ABI3-B3 KKS: R64K, R66K, P69S triple substitution mutation of the ABI3-B3 effector (blue, solid).

#### Evidence for independent origins of LEC2 and FUS3

The B3 domain phylogeny further indicated that LEC2 B3 is more closely related to VP1/ABI3 B3 clade than to the FUS B3 domain. A parsimonious interpretation is that LEC2 and FUS3 diverged independently from VP1/ABI3-type progenitors through loss of the N-terminal COAR domain (Figure 1A and Supplemental Figure S1). The FUS3 clade evidently originated early in the seed plant lineage, whereas LEC2 arose more recently in the rosid clade of angiosperms (flowering plants). Hence, the comparatively recent separation of LEC2 and ABI3 B3 domains afforded an opportunity to identify intrinsic differences in B3 domain structure associated with functional diversification of LEC2 and ABI3 transcription factors.

### A structural basis for functional differentiation of LEC2 and ABI3 B3 domains

#### Identification of clade-specific structural variants

To identify conserved structural features that distinguish of LEC2 and ABI3 B3 domains, variable positions in B3 multiple protein sequence alignments were classified as clade-specific if all amino acid substitutions observed at that position were strictly conserved within each of the four B3 clades (Figure 1B and Supplemental Figure S2). Strikingly, three of four clade-specific amino acid variants that distinguish ABI3 and LEC2 B3 domains are clustered in beta-strand β4 of the protein structure (Figure 1B, C). In B3-DNA complexes (8,9), the C-terminal end of β4 approaches the DNA backbone at an oblique angle (Figure 1C, green ribbon) aligning side-chains of the β4-triad amino acids with the phosphate backbone of DNA.

#### Functional analysis of β4-triad structural variants in yeast

As shown in Figure 1D, the clade-specific β4-triad amino acid substitutions identified by phylogenetic analysis partially account for a qualitative difference in activities of LEC2 and ABI3 B3 domains measured in yeast. To quantify in vivo activities of B3 domains in isolation, we adapted a yeast one hybrid (Y1H) assay (28) based on a minimal *HIS3* promoter containing tandem copies of the Sph/RY cis-element (Figure 1D and Supplemental Figure S3). A GAL4AD:LEC2-B3 effector protein supported growth of transformed yeast cells on selective agar media (Supplemental Figure S3) as well as in liquid culture (Figure 1D). By contrast, a GAL4AD:ABI3-B3 effector did not support growth (Figure 1D). We then analysed a series of reciprocal substitutions that interconvert β4-triad amino acids in the ABI3 and LEC2 B3 domains (Figure 1D). In LEC2-B3 RR, K64R and K66R substitutions combined reduced yeast growth rate compared to wild-type LEC2-B3, whereas the S69P substitution (LEC2-B3 P) by itself completely abolished HIS3 dependent growth. Conversely, introduction of the LEC2 β4-triad variants in ABI3-B3 KKS resulted in a qualitative gain of effector activity (about 60% of LEC2 activity) compared to the inactive wild type ABI3-B3. The P69S substitution (ABI3-B3 S) alone conferred weak activity, whereas combined R64K and R66K substitutions (ABI3-B3 KK) had no effect.

### Differential functions of LEC2 and ABI3 B3 domains measured *in planta*

#### Functional analysis of B3 domains in a complex architecture setting

In developing embryos of plants, protein-protein interactions mediated by the N-terminal COAR domain of VP1/ABI3 proteins physically couple recognition of the Sph/RY cis-element by the C-terminal B3 domain to activities of transcription factors that mediate abscisic acid (ABA) signalling (29–31). To test interoperability of B3 domains from different sources with the COAR domain we used VP1, the maize ortholog of ABI3, as a heterologous domain-swap host for testing B3 function in transgenic Arabidopsis (Figure 2A and Supplemental Figure S4 and S5). As shown previously (23) a ubiquitously expressed *VP1* transgene (*Pro35S:VP1*) 1) complements the green-seed and desiccation-intolerant seed phenotypes of the *abi3-6* null mutant (Supplemental Figure S4), and 2) confers ABA-dependent, ectopic induction of the normally seed specific *Cruciferin-C* (*CRC*) gene in vegetative tissues (Supplemental Figure S5). *CRC* is a well characterized direct target of ABI3 containing multiple Sph/RY motifs in its promoter (32).

**Figure 2.**
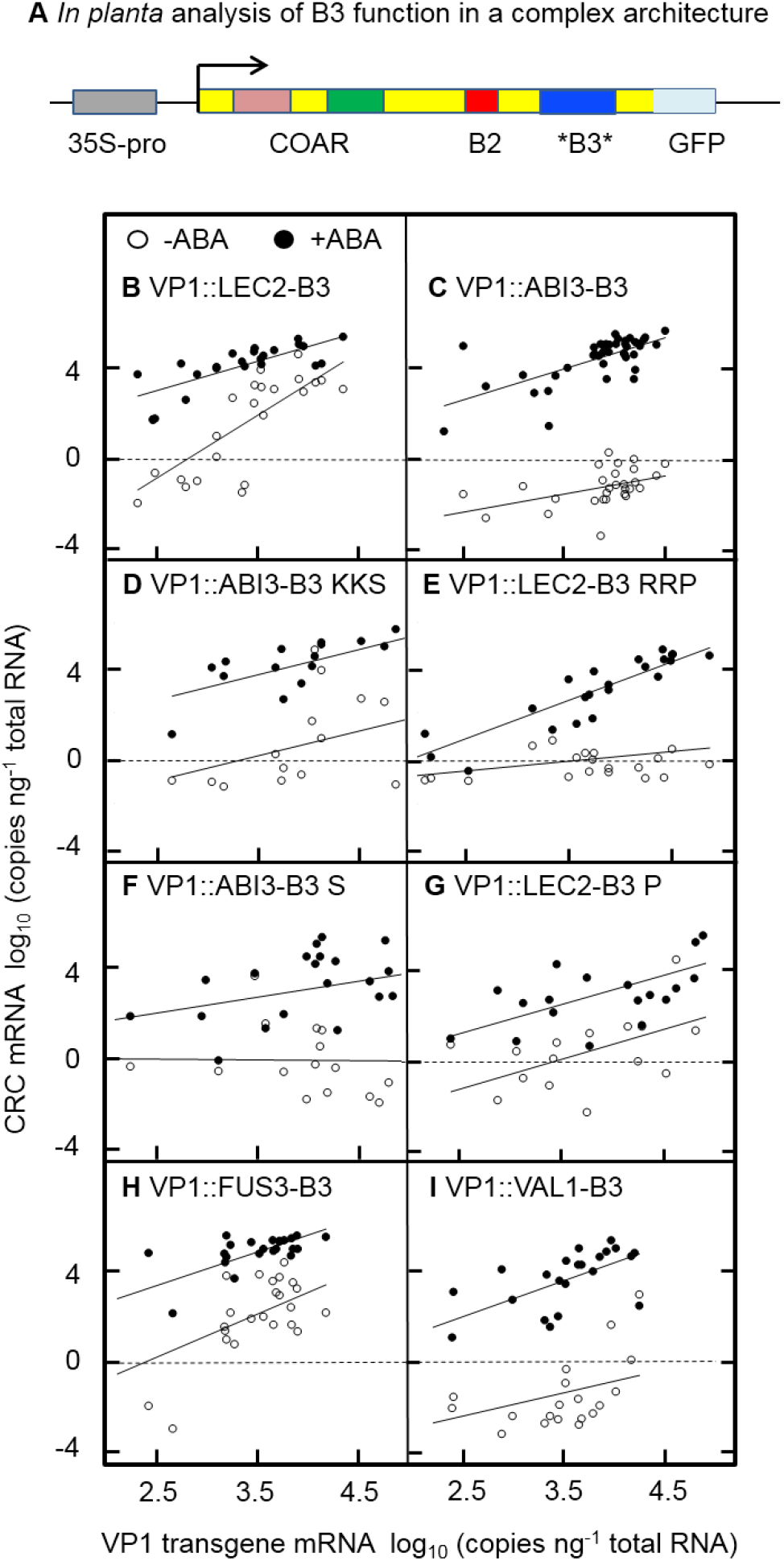
Differential functions of B3 domains in a complex architecture setting. (**A**) To quantify B3 capacities to interoperate with ABA signaling mediated by COAR domain, B3 domain sequences of Arabidopsis ABI3, LEC2, FUS3 and VAL1 proteins were swapped into a heterologous 35S-promoter-VP1 transgene (junctions indicated by *). This transgenic assay takes advantage of the observation that constitutive expression of maize VP1 complements the green embryo and desiccation intolerant seed phenotypes of the Arabidopsis *abi3-6* mutant and conditions ectopic, ABA induction of the endogenous *CRC* gene in leaves (23,33). ABA induction of *CRC* requires both COAR and B3 functions (33). (**B-I**) *VP1::B3* transgene induction of *CRC* with (solid circles) and without ABA treatment (open circles). Expression of the indicated *VP1::B3 transgene* and *CRC* was quantified in individual transformed T1 seedlings by qPCR with and without 24 h treatment with 5 μM ABA. Due to position-effect variation among transformants the assay achieved a >300-fold dynamic range of *VP1::B3* transgene expression. (**B, C**) In contrast to *VP1::LEC2-B3*, induction of *CRC* by *VP1::ABI3-B3* shows strict dependence on ABA signaling. (**D, E**) Reciprocal β4-triad substitutions in ABI3-B3 and LEC2-B3 domains interchange capacities for ABA-independent induction of *CRC* (open circles). (**F, G**) Reciprocal serine-proline substitutions at position 69 of LEC2 and ABI3 B3 domains have intermediate effects on ABA-dependent and -independent transgene induction of *CRC*. (**H, I**) Functional similarities of FUS3 B3 and VAL1 B3 domains to LEC2 B3 and ABI3 B3 domains, respectively, in the transgenic assay.

Crucially, ABA-dependent induction of *CRC* in vegetative tissues requires both COAR and B3 domains of VP1, whereas the COAR domain alone is sufficient for rescue of *abi3-6* seed maturation phenotypes (33). As a measure of B3 activity, we quantified expression of both the *VP1::B3* transgene and endogenous *CRC* gene in detached leaves of individual 14- day-old T1 transgenic seedlings using qPCR following 24h incubation on MS media with or without a 5μM ABA treatment (Figure 2 and Supplemental Figure S5). To confirm that ectopic induction of *CRC* in the transgenic assay required B3 domain activity (33), K15R loss-of-function mutations were tested for each B3 domain as negative controls (Supplemental Figure S6). Due to position effect variation among individual transformants, the analysis spanned a >300-fold dynamic range of transgene expression (~2.5 log units, Figure 2B, C, x-axes).

#### β4-triad differentiation of B3 domain interoperability with the COAR domain

As shown in Figure 2B, C, *abi3-6* seedlings transformed with either *VP1::LEC2-B3* or *VP1::ABI3-B3* exhibited ABA dependent induction of *CRC* over a wide range of transgene dosage. By contrast, the VP1::LEC2-B3 and VP1::ABI3-B3 proteins show strikingly different capacities for induction of *CRC* in the absence of ABA (Figure 2B, C, open circles).

To determine whether this qualitative difference in interaction with ABA signalling was attributable to the β4-triad structural variant, we tested effects of reciprocal β4-triad substitutions in VP1::LEC2-B3 and VP1::ABI3-B3 proteins (Figure 2D-G). The LEC2-B3 RRP triple substitution nearly abolished ABA-independent *CRC* activation (Figure 2E, open circles) while retaining a strong capacity for ABA dependent induction of *CRC* (Figure 2E, solid circles). Thus, in key respects the LEC2-B3 RRP triple substitution was functionally equivalent to wild type ABI3-B3 (Figure 2C). Conversely, the ABI3-B3 KKS β4-triad substitution exhibited a marked capacity for ABA independent activation of *CRC* in seedlings (Figure 2D, open circles), a response resembling activity of wild type LEC2-B3 (Figure 2B). Consistent with the Y1H results, the effects of reciprocal single amino acid substitutions at position 69, ABI3-B3 S (Figure 2F) and LEC3-B3 P (Figure 2G) were intermediate albeit with markedly increased variation in transgene dosage and ABA responses. Hence, the β4-triad partially accounts for differential in vivo activities of LEC2 and ABI3 B3 domains both in yeast and *in planta*.

We reasoned that enhanced activity of LEC2 B3 domain relative to the ABI3 B3 domain is a plausible adaptation of LEC2 B3 to a reduced protein architecture. In the B3 network, LEC2 functions as a pioneer transcription factor that interacts primarily with the Sph/RY ciselement (14,15). By contrast, ABI3 B3 activity is constrained to interoperate with the COAR domain which is capable of independently tethering ABI3 to target promoters via interaction with ABA regulated bZIP factors (30). By tethering ABI3 to the *CRC* promoter, ABA regulated bZIP transcription factors increase the local concentration of B3 in vicinity of Sph/RY lowering the DNA-binding affinity that is optimal for a conditional interaction between B3 and COAR domains. In the same architecture, hyper-active DNA binding activity of the LEC2 B3 domain causes transcriptional activation in the absence of COAR tethering thereby superseding proper integration of ABA-signalling.

#### Extension of the architecture complexity hypothesis to FUS3 and VAL1 B3 domains

To test the architecture complexity hypothesis, we determined whether *in planta* activities of FUS3-B3 (Figure 2H) and VAL1-B3 (Figure 2I) domains show a similar correlation with architectural complexity. As noted above, FUS3 likely evolved a reduced domain architecture independently of LEC2. Although VAL1 and ABI3 have only the B3 domain in common, analogous to the ABI3, the complex architecture of the VAL1 repressor serves to physically integrate B3 activity with domains that independently recognize chromatin marks (3,6,7,12,13). In the VP1::B3 transgenic assay, activity of FUS3-B3 (Figure 2H) is remarkably similar to LEC2-B3 (Figure 2B) in conferring strong ABA independent induction of *CRC*, whereas VAL1-B3 (Figure 2I) closely resembles ABI3-B3 in lacking capacity for ABA independent activation of *CRC* except in seedlings with very high transgene dosages.

#### Novel developmental phenotypes associated with hyper-active LEC2 and FUS3 B3 domains

The functional dichotomy between LEC2/FUS3 and ABI3/VAL1 B3 domains is further revealed at the level of phenotype. A subset of *VP1::FUS3-B3* and *VP1::LEC2-B3* transformants produced abnormal seedlings with features resembling embryonic phenotypes associated with *val* loss of function mutants (4,5) (Figure 3A, Supplemental Figure S7 and Supplemental Table S1). By contrast, embryonic seedling phenotypes were rare among VP1::ABI3-B3 and VP1::VAL1-B3 transformants (Figure 3B). In *val* loss-of-function mutants, embryonic seedling phenotypes are associated with up-regulation of *LEC1*, a transcription factor implicated in initiation of embryogenesis (4,5). Ectopic activation of *LEC1* was evident in all *VP1::FUS3-B3* and *VP1::LEC2-B3* transformants that developed embryonic features (Figure 3C, open circles) as well as in a majority of transformants with normal seedling phenotypes (Figure 3C, solid circles) indicating that induction of *LEC1* was necessary but not sufficient for induction of embryonic seedling phenotypes. In line with that conclusion, *LEC1* expression in *VP1::ABI3-B3* and *VP1::VAL1-B3* transformants was consistently low. Moreover, *VP1::LEC2-B3* transformants with S69P and RRP triple substitution showed reduced *LEC1* induction, whereas slight induction of *LEC1* was conferred by *VP1::AB3-B3 S* and *KKS* substitutions (Supplemental Figure S8).

**Figure 3.**
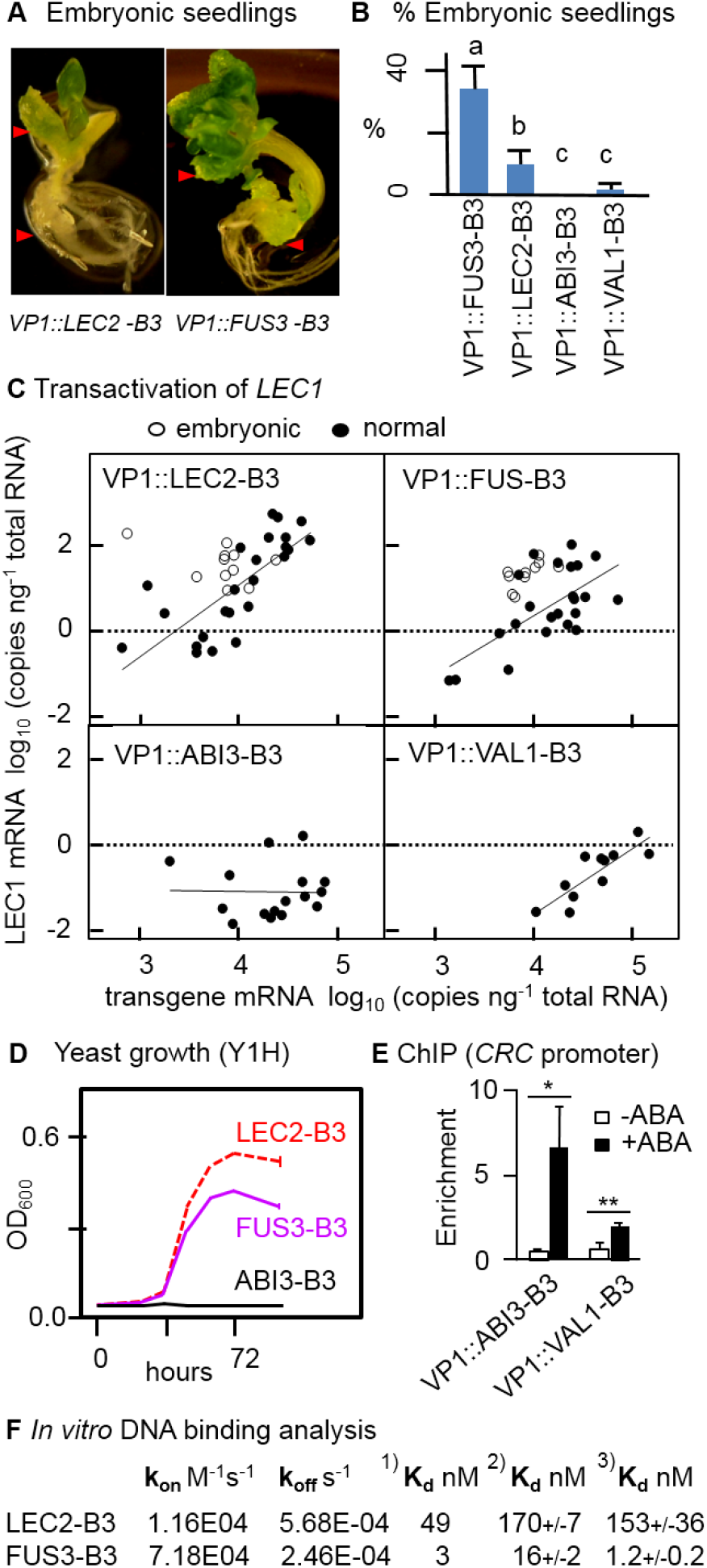
A functional dichotomy of B3 domains associated with complex and reduced architectures. (**A**) Embryonic seedling phenotypes induced by *VP1::LEC2-B3* and *VP1::FUS3-B3* transgenes. (**B**) Proportions of transformants with embryonic seedling phenotypes (Supplemental Table S1). Error bars are standard error of the mean (for n=3 or n=4 independent transformation experiments). Values with different letters are significantly different at p<0.05 based on pair-wise t-tests. (**C**) Induction of embryonic seedling phenotypes by *VP1::LEC2-B3* and *VP1::FUS3-B3* transgenes is associated with capacity for ectopic activation of *LEC1*. Expression of *LEC1* and *VP1::B3* transgenes was quantified by qPCR in T1 transformants with normal phenotypes (solid circles) and embryonic phenotypes (open circles). Seedlings were grown on MS media without ABA. (**D**) FUS3-B3 supports growth of yeast in the Y1H assay. Y1H analysis was performed as described in Figure 1D and Supplemental Figure S3. (**E**) ChIP-qPCR assays confirm ABA-dependent binding of *VP1::ABI3-B3* and *VP1::VAL1-B3* proteins to the *CRC* promoter. Error bars are standard error of the mean (n=3) (unpaired t test; * denotes P<0.05, ** denotes P<0.005). (**F**) *In vitro* DNA binding activities of FUS3 and LEC2 B3 domains. *In vitro* binding of purified recombinant GST-B3 protein to an oligonucleotide containing an Sph/RY dimer was performed using the Octet instrument (26) and by EMSA (22). Values for kon and koff were determined from the Octet kinetic analysis using Octet Data Analysis HT software. K_d_ values were estimated using three methods: 1) ratio of k_off_ and k_on_; 2) Octet equilibrium analysis, and 3) EMSA densitometry measurements fit to the Hill equation in R (Supplemental Figure S12). Estimates for the Hill constants of LEC2 and FUS3 B3 domains did not differ significantly from 1.0.

Other properties of FUS3 B3 and VAL1 B3 domains fit the architecture complexity hypothesis. Like LEC2 B3, the FUS3 B3 domain is highly active in the Y1H assay (Figure 3D) under conditions in which VAL1 B3 and ABI3 B3 are in-active (Supplemental Figure S3). Although VAL1 and ABI3 have very different architectures, the similar capacities of VP1::VAL1-B3 and VP1::ABI3-B3 proteins to support ABA dependent induction of *CRC* suggests functional equivalence in the VP1 setting. ChIP analyses confirm that the VP1::VAL1 B3 transcription factor faithfully recognizes the *CRC* promoter in an ABA dependent manner, albeit with less efficiency than VP1::ABI3-B3 (Figure 3E). Interestingly, functionally similar VAL1 B3 and ABI3 B3 domains share two of three β4-triad variants (Figure 1B).

#### Functional differences of LEC2 and FUS3 B3 domains

Although LEC2 and FUS3 B3 domains have qualitatively similar functionality in yeast and *in planta* assays there are striking quantitative differences (Figure 3B). In DNA binding assays recombinant FUS3 B3 protein exhibits 10-to 100-fold greater affinity for Sph/RY sequence in vitro compared to LEC2 B3 (Figure 3F, Supplemental Figure S9-12). K_d_ estimates for FUS3 B3 based on three different methods ranged from 1.2 nM to 16.0 nM compared to a range of 49.0 nM to 170.0 nM for the K_d_ of LEC3 B3. Kinetic analysis indicates that the lower K_d_ of FUS3 B3 is partly attributed to a ~6-fold faster “on” rate compared to LEC2 B3. Conceivably, the higher binding affinity of FUS3 B3 would account for the 3-fold greater frequency of embryonic seedling phenotypes observed in *VP1::FUS3-B3* transformants compared to *VP1::LEC2-B3* transformants (Figure 3B). Under identical *in vitro* conditions, very high concentrations of protein required for detection of DNA binding by ABI3 B3 (Supplemental Figure S13A) and VAL1 B3, which failed to bind even at high concentrations, were not conducive to determination of K_d_ values (Supplemental Figure S13B).

### DNA backbone interactions are a structural basis for B3 adaptation to architecture

#### Rationalizing effects of the β4-triad on B3 domain DNA binding activity

To rationalize the structural implications of the β4-triad in differentiation of LEC2 and ABI3 B3 domains, we analysed published structures of B3 domains bound to the cognate Sph/RY element (8,9) and constructed homology models of ABI3 wild type and mutant B3 domains using VAL1 and FUS3 structures (PDB id: 6J9A and 6J9B) (9) as templates. We noticed that in the LEC2 B3-DNA complex, the peptide backbone of β4 is positioned closer to the phosphate backbone of DNA than in the VAL1 B3-DNA complex (Figure 4A-C) with the point of closest approach occurring at β4-triad residue 69. In the LEC2 B3-DNA complex, a 2.5 Å reduction in distance of the peptide backbone from the nearest DNA phosphate (3.2 Å in LEC2 vs 5.75 Å in VAL1, Figure 4D-E; and 4.67 Å in FUS3, Figure 4F) is accommodated by formation of a H-bond between the amino nitrogen of S69 and a phosphate of the DNA backbone (Figure 4G). By contrast, the corresponding nitrogen of proline 69 in VAL1 and ABI3 lacks capacity for H-bonding suggesting that H-bond formation accounts for the pronounced impact of the S69P substitution on B3 domain activity (Figure 4H-K).

**Figure 4.**
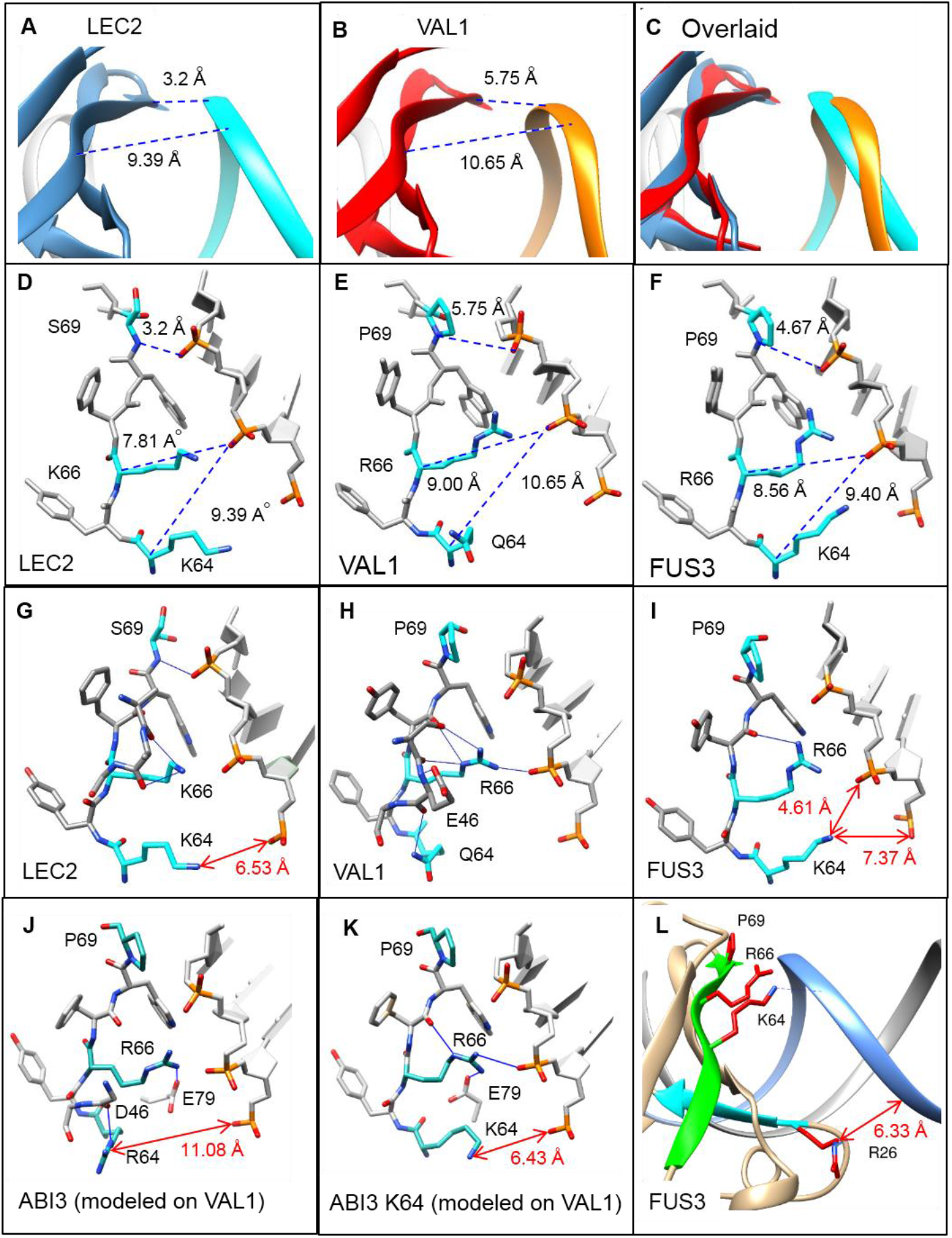
Structural basis for functional differentiation of B3 domains. (**A, B**) Distance separating β4 from the phosphate backbone of DNA in LEC2 (PDB ID: 6J9C) and VAL1 B3 (PDB ID: 6J9A) structures. Dashed lines indicate distance between the peptide backbone at position 64 (alpha carbon) and 69 (amine nitrogen) and nearest phosphate oxygen atom in the DNA backbone. Distances were measured in angstroms using Chimera (18). (**C**) Overlay of LEC2 (protein blue, DNA backbone light blue) and VAL1 (protein red, DNA backbone gold) structures (9). (**D-F**) Detailed view of backbone separations in LEC2 (PDB ID: 6J9C), VAL1 (PDB ID: 6J9A) and FUS3 (PDB ID: 6J9B) B3 domains. β4-triad amino acid side chains are colored blue. Distances (dashed lines) were measured in angstroms using Chimera (18). (**G-K**) H-bonds formed by β4-triad amino acids in B3 domains. H-bonds (blue lines) were identified using the Chimera FindHBond tool with relaxed constraints (18). Red arrows and text indicate distance of positively charged atoms of amino acid 64 from phosphates of the DNA backbone. Models of ABI3 WT and ABI3 K64 mutant structures in J) and K) were constructed in MODELLER (17) using VAL1 (PDB ID: 6J9A) (9) as template. (**L**) Position of the arginine 26 side-chain in FUS3 B3 consistent with a weak electrostatic interaction with the DNA backbone. The distance of the guanidinium side-chain from the nearest phosphate is indicated with a red arrow and text.

In all B3 structures (9), the positively charged side-chain of residue 66 interacts electrostatically with the DNA backbone. In the LEC2 B3-DNA complex reduced separation of the peptide and phosphate backbones at position 66 (7.81 Å in LEC2 vs 9.0 Å in VAL1) is accommodated by the shorter side-chain of the lysine compared to arginine 66 in VAL1. Hence, we suggest that the R66 side-chain in ABI3 limits proximity of the β4 backbone to the phosphate backbone. An intermediate backbone separation observed in the FUS3 B3 structure (8.56 Å, Figure 4F) is associated with a retracted conformation of the R66 sidechain relative to VAL1 R66 (Figure 4E).

Finally, residue 64 is known to have an important, albeit enigmatic role in B3 interaction with DNA (8,9). Interestingly, in ABI3 B3 domain models (Figure 4J) based on independent VAL1 B3 structures (8,9) as well as FUS3 B3 (9) (Supplemental Figure S14A and S14B), the R64 side-chain consistently formed a H-bond with the backbone carbonyl of amino acid D46. While the amide nitrogen of the Q64 side-chain in VAL1 forms an analogous H-bond (Figure 4H), the K64 side-chains of LEC2 (Figure 4G) and FUS3 (Figure 4I) do not, favouring extension of the lysine side-chain toward phosphate backbone. Loss of the H-bond at position 46 in B3 domains with the K64 substitution can be rationalized – because in addition to being shorter, the lysine amine typically has less facility for H-bonding than the guanidinium of arginine. Consistent with that interpretation, loss of the D46 H-bond in a model of the ABI3 B3 K64 mutant (Figure 4K) is associated with a pronounced shift of the amino acid 64 side-chain toward the phosphate backbone (11.08 Å in WT ABI3 compared to 6.43 Å in the K64 mutant). The comparable distance of the K64 side-chain from the phosphate backbones in LEC2 (6.53 Å) and FUS3 (4.61 Å) is consistent with a weak electrostatic interaction with DNA that is not present in DNA bound VAL1 and ABI3 B3 domains.

#### Variation in non-specific DNA backbone interactions that preserves sequence specificity

Together these observations point to subtle variation in the interaction of β4 with the DNA backbone as an underlying structural basis for functional differentiation of LEC2 and VP1/ABI3 type B3 domains. Importantly, β4-triad variants have little apparent effect on base contacts that determine sequence specificity of B3 domain binding to DNA. Hence, variation affecting a non-specific electrostatic contribution to the protein-DNA interaction provides a mechanism for differentially altering DNA binding activity and/or affinity of B3 domains while preserving DNA sequence specificity. Because a non-sequence-specific electrostatic contribution to the total binding free energy is a universal feature of protein-DNA interactions (34,35), this mechanism has broad relevance to functional diversification of transcription factor families. In this context, we suggest that architectural constraints are a key driver for functional differentiation of conserved DNA binding domains.

The B3 network is an attractive model for further exploration of the relationship between architecture and DNA binding domain evolution. Intriguingly, FUS3 B3 and LEC2 B3 domains likely derived independently from ABI3-like ancestors (Figure 1A and Supplemental Figure S1). FUS3 B3 has only one β4-triad variant (K64) in common with LEC2 suggesting that its high DNA binding affinity and enhanced capacity for induction of embryonic development has a different structural basis. Hence, the functional similarities that distinguish them from ABI3 and VAL1 B3 domains are evidently due to convergent evolution. We note that a second cluster of clade-specific variants near β2 includes a glycine to arginine substitution at position 26 that is unique to the FUS3 B3 domain (Figure 1B). Interestingly, positioning of the R26 side-chain in the FUS3 B3 structure is consistent with a weak electrostatic interaction (6.53 Å separation, Figure 4L) with a phosphate of the same DNA strand engaged by the β4 triad. While the functional significance of this clade-specific variant remains to be tested, the R26 substitution fits the broader theme of subtle variation in electrostatic interactions playing a central role in functional diversification of B3 domains.

The precise roles of comparatively weak electrostatic interactions and separation of β4 from the DNA backbone in determining binding affinity are not clear. A deeper understanding may result from insight into effects of β4-triad substitutions on the dynamics of B3 domain binding to DNA. Dynamic properties are likely to be especially important *in vivo* where transcription factors must efficiently scan the genome for cognate binding sites in the nucleus.

## ACCESSION NUMBERS

Sequence data from this article can be found in the Arabidopsis Information Resource (TAIR) and GenBank/EMBL databases under the following accession numbers: ABI3, TAIR AT3G24650; FUS3, TAIR AT3G26790; LEC2, TAIR AT1G28300; VAL1, TAIR AT2G30470; VAL2, TAIR AT4G32010; VAL3, TAIR AT4G21550; LEC1, TAIR AT1G21970; CRC, TAIR AT4G28520; TUB2, TAIR AT5G62690; and VP1, GenBank NM_001112070.

## SUPPLEMENTARY DATA

Supplementary Data are available at NAR online.

## ACKNOWLEDGEMENT

The authors are grateful for the expert technical assistance of Shan Wu and support from University of Florida Institute of Food and Agriculture Sciences. We Thank Eiji Nambara (University of Toronto) for providing the *abi3-6* stocks and Dr. Karen Koch and her lab for advice and access to real-time PCR equipment.

---

**Supplemental Figure S1.**
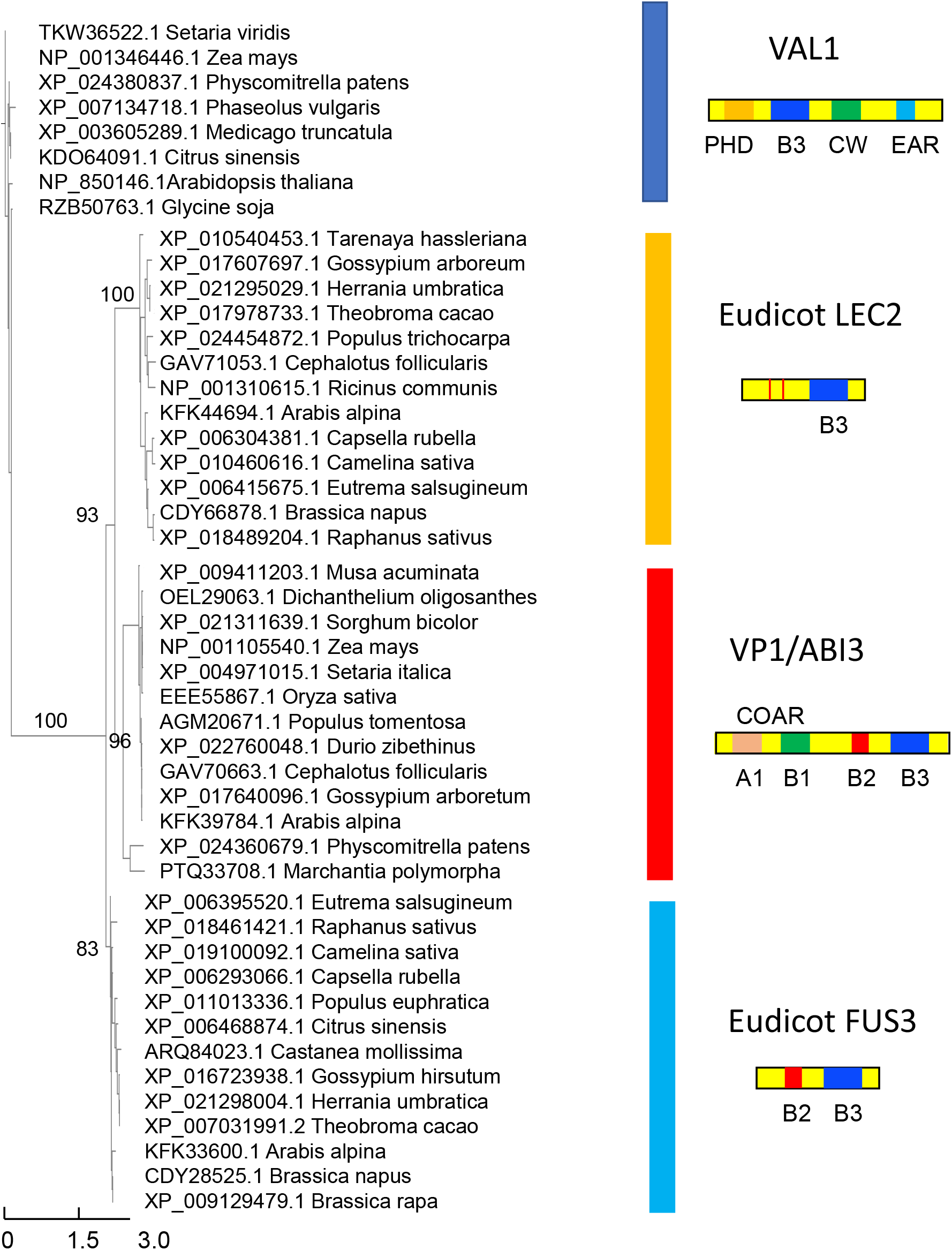
Phylogenetic tree of AFL and VAL B3 domain sequences. B3 domain sequences (120 amino acids) were aligned using MAFFT (19) and a maximum likelihood tree constructed in PhyML (20) with bootstrap support based on BOOSTER (21). Domain architectures are diagrammed for each of the four principle B3 clades, VAL1, dark blue rectangle, LEC2, gold rectangle, VP1/ABI3 red rectangle, FUS3, light blue rectangle..

**Supplemental Figure S2.**
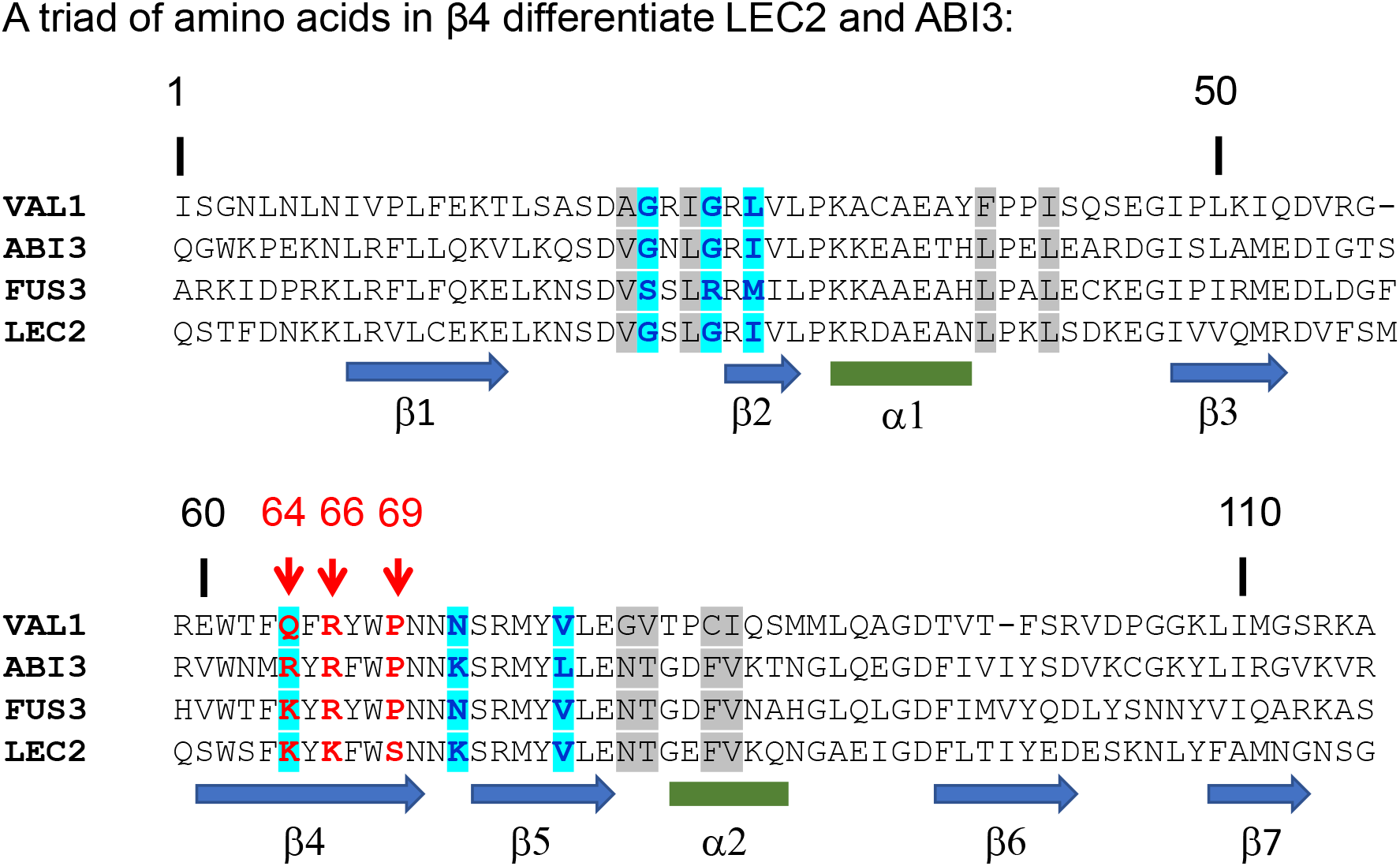
Alignment of Arabidopsis AFL and VAL1 B3 domains. Clade-specific variant positions are highlighted as described in Figure 1B. β4-triad positions are colored red and indicated by arrows. Beta-sheet (block arrows) and helix regions (rectangles) are based on a consensus of B3 structures.

**Supplemental Figure S3.**
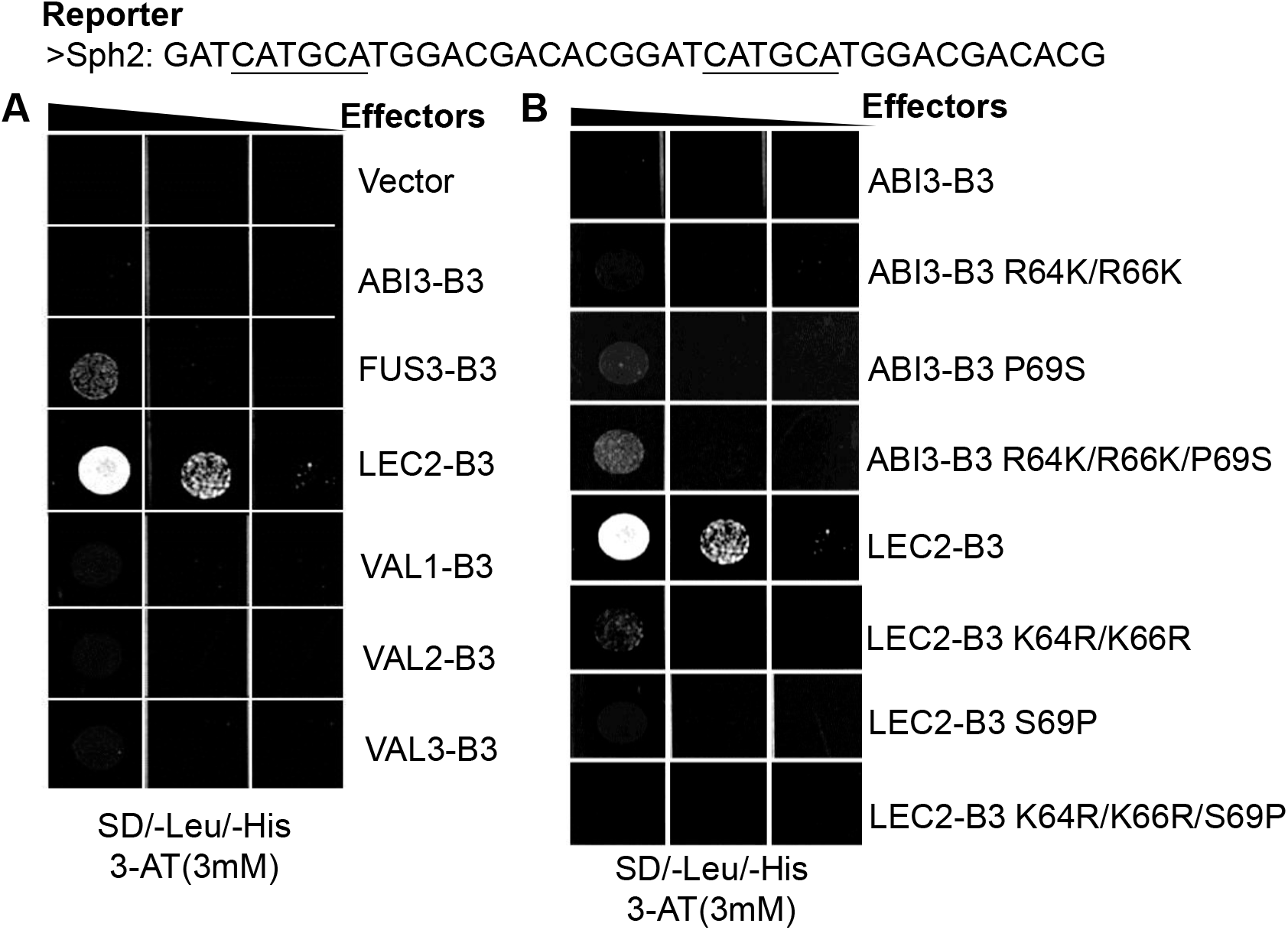
Functional analysis of B3 domains in a Y1H assay showing partial interconversion of LEC2 and ABI3 B3 domain activities. AFL and VAL B3 domain sequences used in the effector constructs are shown in Supplemental Figure S2. The Y1H system: yeast reporter strains carry the *HIS3* gene under control of a *Sph2* minimal promoter. The core Sph motifs are underlined in the Sph2 minimal reporter sequence. All Y1H effectors were fusions to a GAL4 activation domain in the pGADT7 vector, which contains a *LEU2* marker for selection in yeast. (**A, B**), Colony growth assays of yeast transformed B3 domain effectors from either AFL or VAL. Equal volumes of ten-fold serial dilutions of effector transformed yeast cells were spotted on SD/-Leu/-His plus 3mM 3-AT media, and incubated at 30°C for 4 days. (**A**) AFL and VAL WT B3 domain effectors; (**B**) LEC2 B3 and ABI3 B3 domain mutant effectors.

**Supplemental Figure S4.**
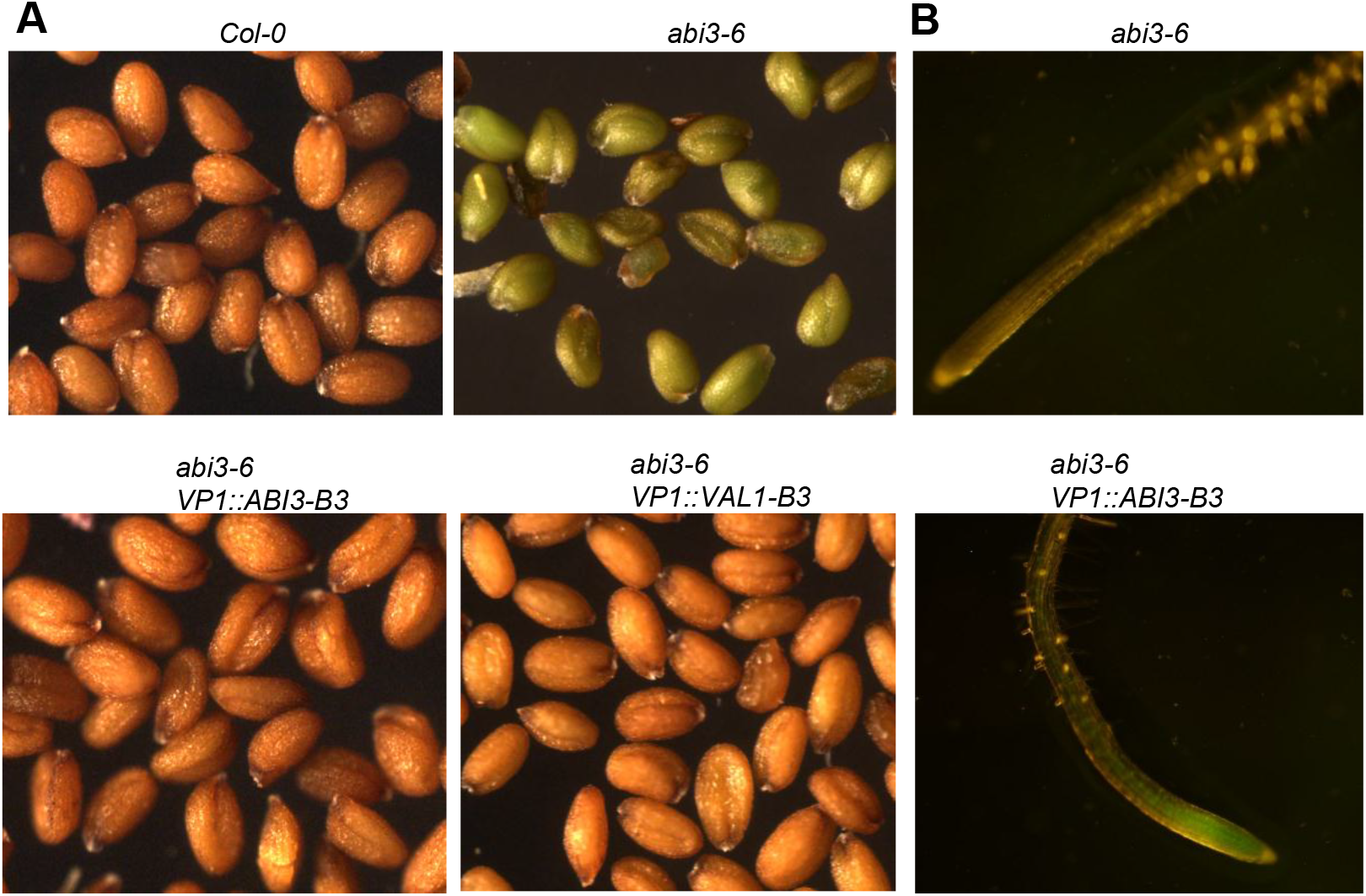
The seed phenotype and GFP expression of the *Pro35S:VP1::B3* transgenic plants. (**A**) *Pro35S:VP1::ABI3-B3* and *Pro35S:VP1::VAL1-B3* complemented *abi3-6* mutant seed green color and desiccation intolerance phenotypes. Mature siliques were collected from Columbia (*Col-0*) wild type, *abi3-6, abi3-6 Pro35S:VP1::ABI3-B3* and from *abi3-6 Pro35S:VP1::VAL1-B3* T3 homozygous transgenic plants. (**B**) GFP expression was observed in root tips of 12-d-old *abi3-6 Pro35S:VP1::ABI3-B3* T3 homozygous transgenic plants

**Supplemental Figure S5.**
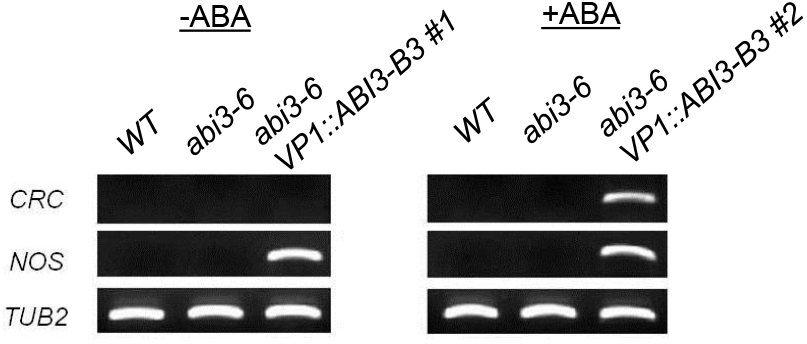
*CRC* expression in the leaf tissue of *WT, abi3-6*, and the T1 line of *abi3-6 Pro35S:VP1::ABI3-B3* with and without ABA treatment. RNA was isolated from leaves of 14-d-old T1 transgenic seedlings in absence or in presence of ABA treatment. For this experiment, 2 independent T1 *abi3-6 Pro35S:VP1::ABI3-B3* seedlings that had normal phenotypes were used. RT-PCR was performed to detect the transcript of downstream target *CRC* and transgene. Primers targeted to the NOS-terminator region were used for quantify the transcript level of the *VP1::ABI3-B3* transgene. The amount of total RNA used for each reaction is 25ng. *TUB2* was used as an endogenous control.

**Supplemental Figure S6.**
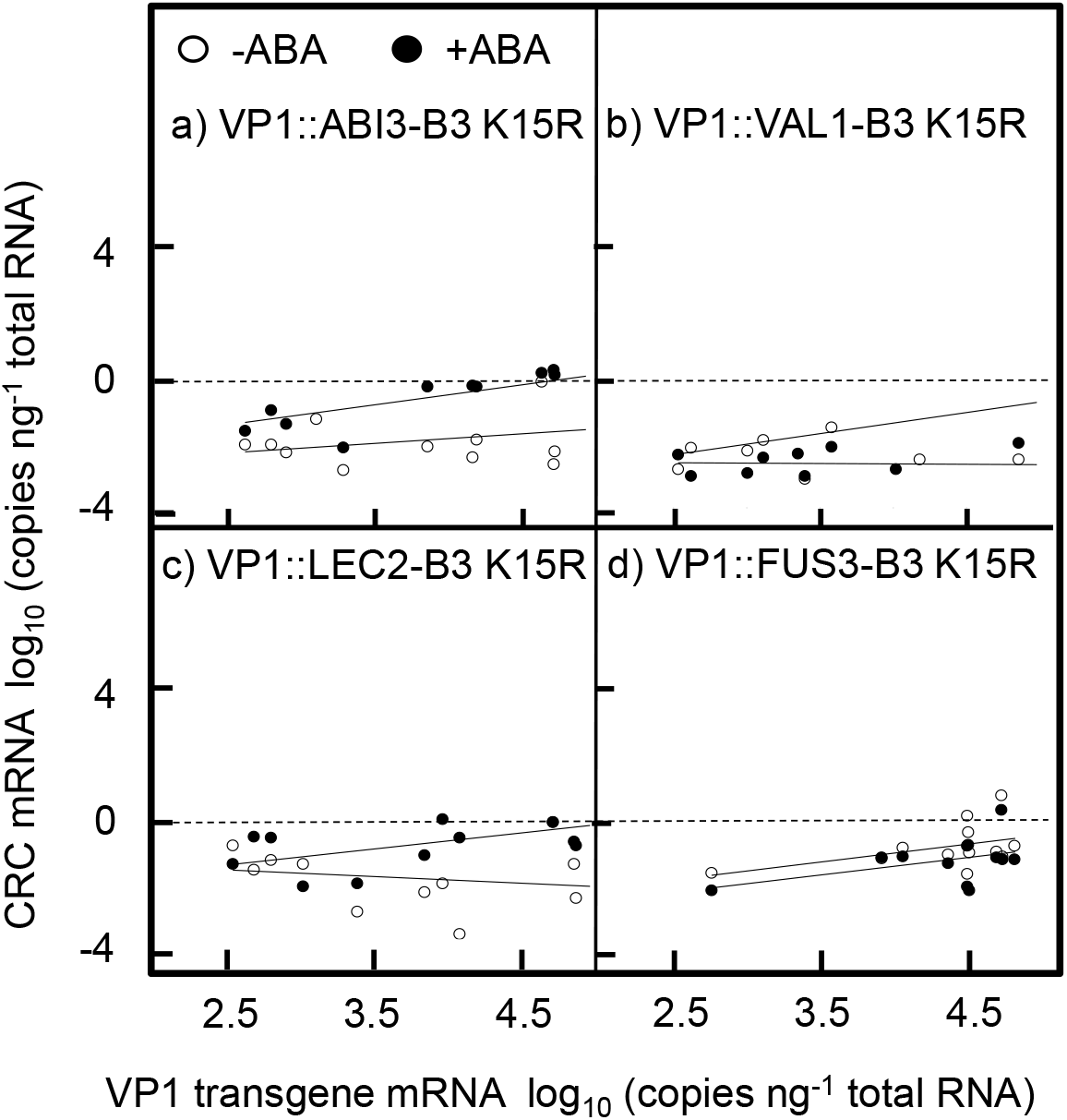
The K15R loss of function mutant^15^ abolishes *in vivo* activities of ABI3, LEC2, FUS3, and VAL1 B3 domains. Expression of the indicated *VP1::B3* transgene and *CRC* was quantified in individual transformed T1 seedlings by qPCR with and without 24 h treatment with 5 μM ABA.

**Supplemental Figure S7.**
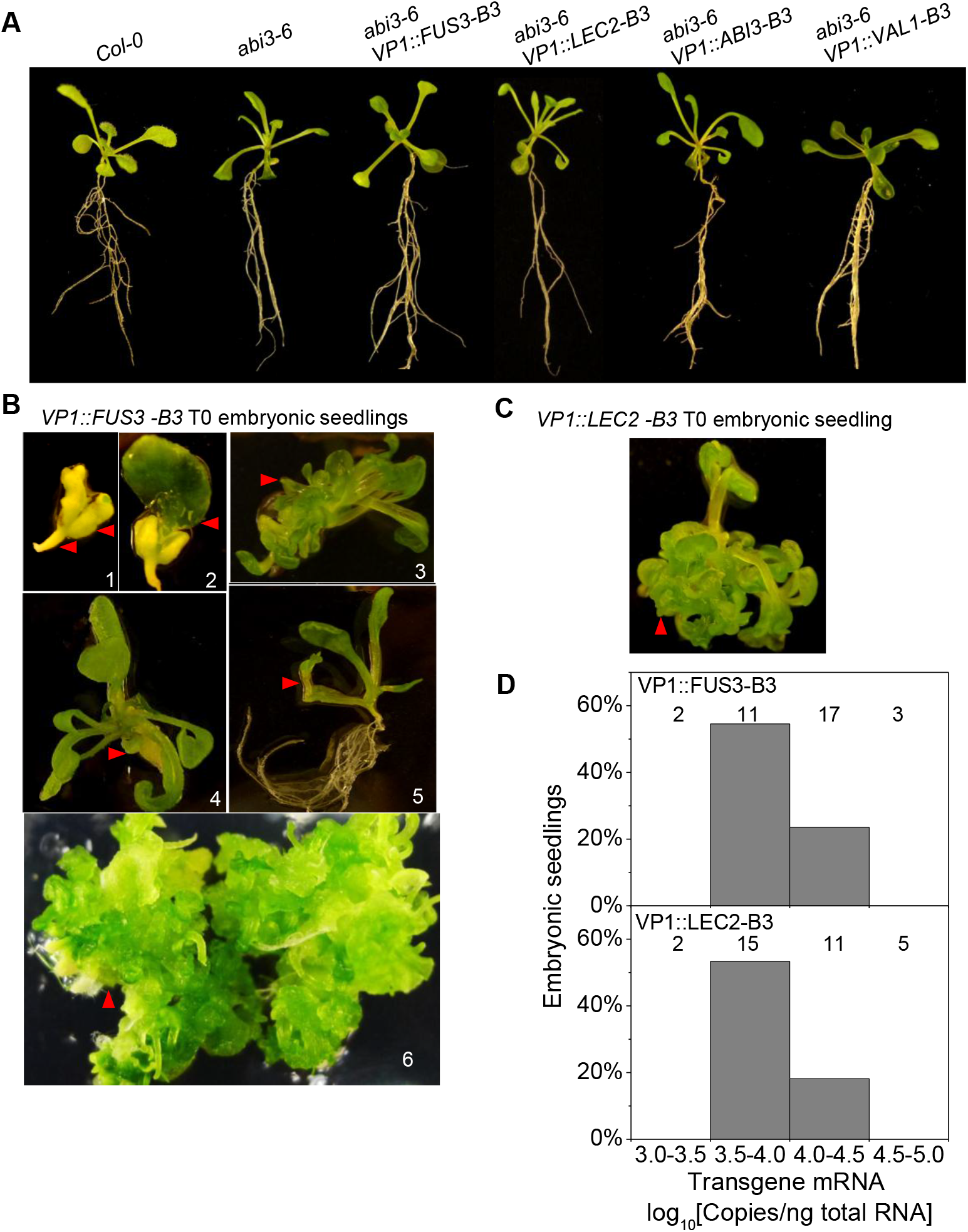
Occurence of embryonic seedling phenotypes among *P35S:VP1::LEC2-B3* and *P35S:VP1::FUS3-B3* transformants. *Pro35S: VP1::B3* chimeral transgenes were used to transform to *abi3-6* mutant Arabidopsis plants. T1 transformants that complemented the desiccation intolerant seed phenotype of *abi3-6* were grown 14-d on MS media containing Hygromycin. (**A**) T1 transgenic seedlings with normal phenotypes compared to *WT* and *abi3-6* seedlings. (**B**) The *abi3-6 Pro35S:VP1::FUS3-B3* T1 transgenic seedlings having a variety of embryonic seedling phenotypes involving callus formation (red arrows). 1, a seedling with arrested embryos; 2, a seedling with arrested embryos that produced a shoot from the SAM; 3, a seedling with many adventitious shoots but no primary root; 4, a seedling with near-normal shoot development, but callus formation in the root region; 5, a seedling with callus in the cotyledon region; 6, an embryonic seedling with extensive callus tissue as well as adventitious shoot and root formation after one month culture on MS media. (**C**) Representative *abi3-6 Pro35S:VP1::LEC2-B3* T1 transgenic seedling that have embryonic seedling phenotype. (**D**) Histogram of the frequency distribution of embryonic seedlings. Frequency distribution of embryonic seedlings classified based on transgene expression in *abi3-6 Pro35S:VP1::FUS3-B3* and *abi3-6 Pro35S:VP1::LEC2-B3* transgenic lines. The total number of seedlings in each class is indicated above the bar.

**Supplemental Figure S8.**
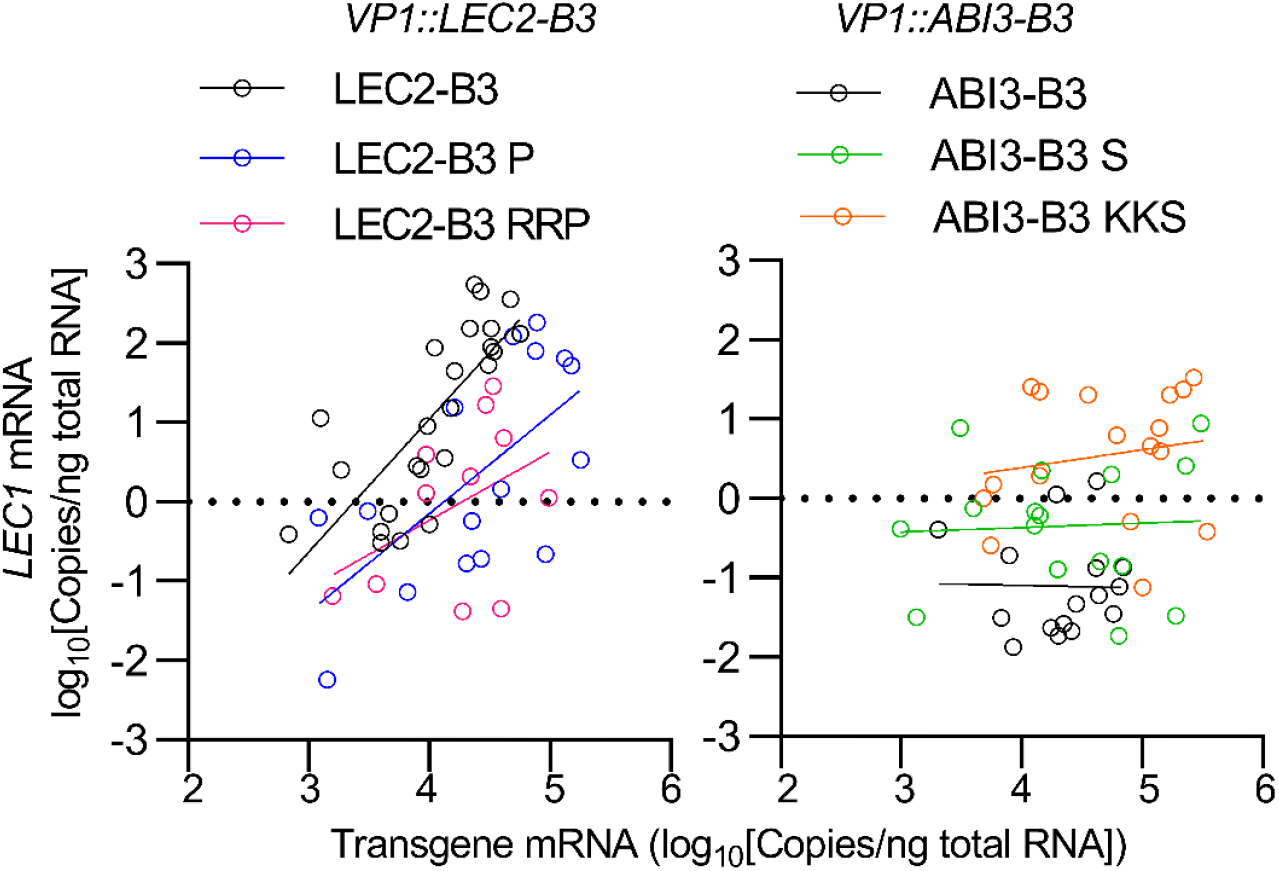
*LEC1* expression in the *P35S:VP1::LEC2-B3* and *P35S:VP1::ABI3-B3* transformants. 14-d-old T1 *Pro35S:VP1::B3* transgenic seedlings were analyzed for transgene and *LEC1* expression using qPCR. Total RNA was isolated from leaf tissue of normal seedlings.

**Supplemental Figure S9.**
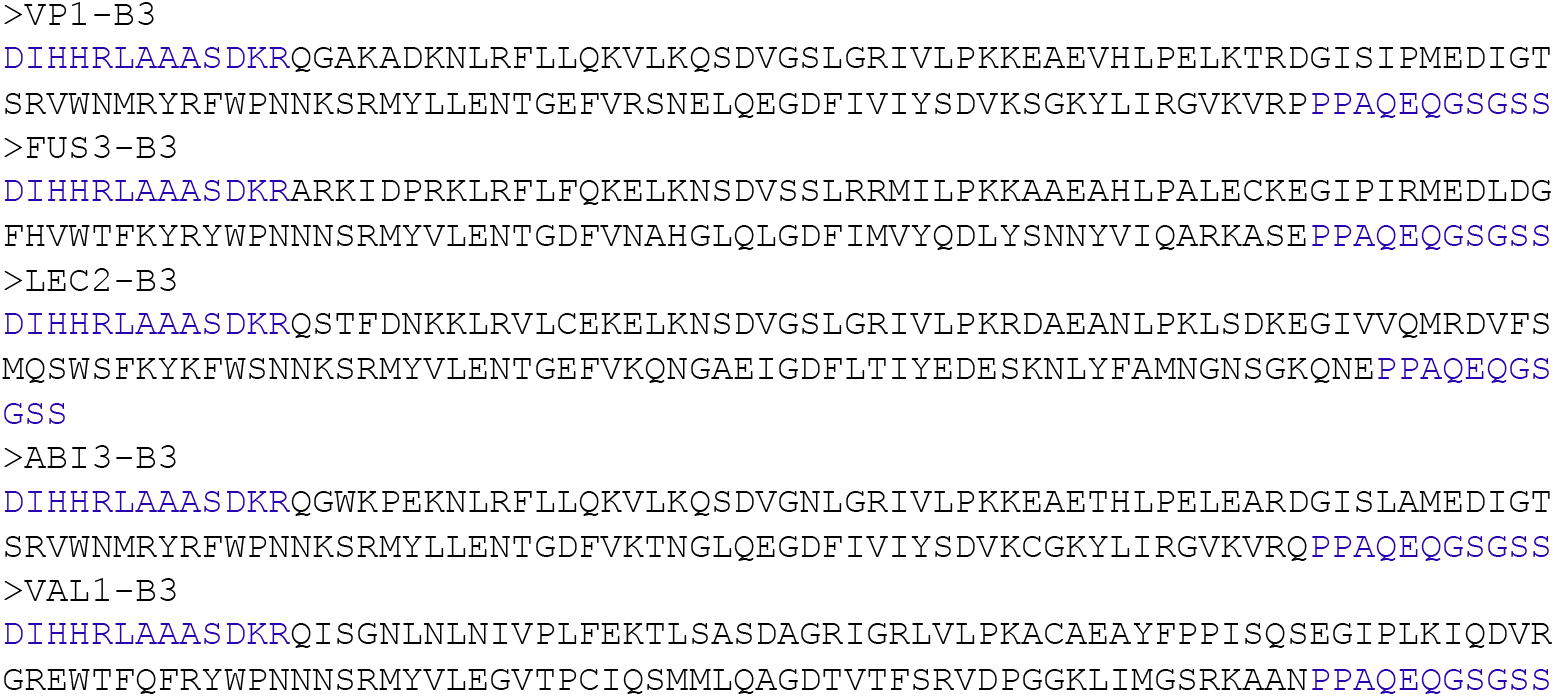
Amino acid sequences of B3 domains of GST-B3 recombinant proteins used in the *in vitro* binding assays. VP1-B3 sequence contains 140 amino acid residues was used previously in in vitro DNA binding analysis^26^. The AFL and VAL B3 domain residues in black font were used in Y1H and also used for making all the domain replacement in the *VP1::B3* chimeral transgenes (Figure 2A). The blue residues of N and C terminal of VP1 B3 domain are contained in all the AFL and VAL1 B3 domains tested in the *in vitro* binding assays.

**Supplemental Figure S10.**
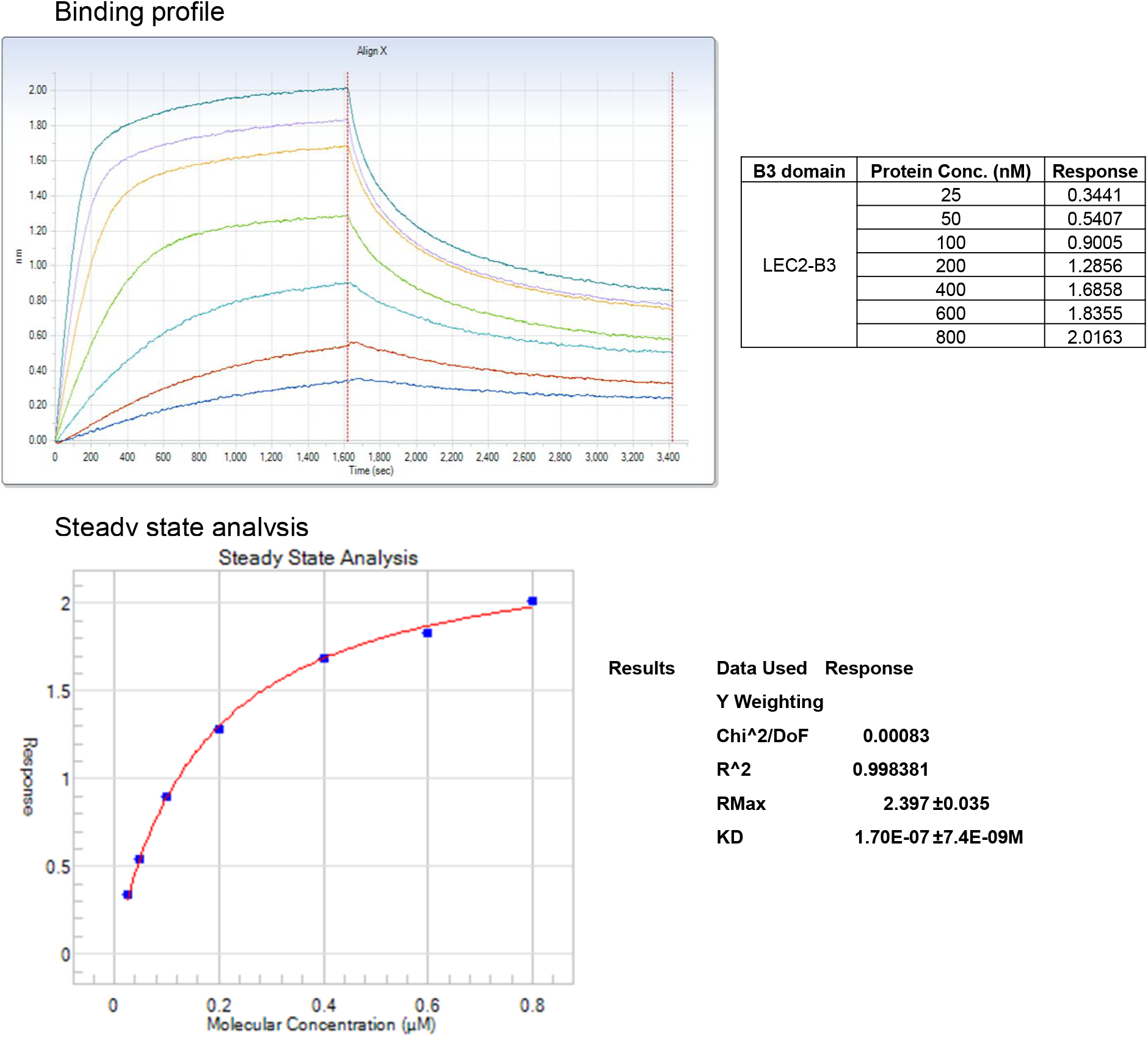
Quantification of LEC2-B3 domain DNA binding activity by Octet Qke. B3-DNA-binding was measured in a label-free in vitro kinetics assay at pH 7.0 using Octet Qke system (Pall ForteBio). Detail experimental conditions and procedures are described in Methods. Measurements were taken from one experimental replicate. The association rate (k_on_), dissociation rate (k_off_), kinetic K_d_ and steady state K_d_ were obtained by Octet Data Analysis HT software fitting a one-site binding curve using non-linear regression. Similar results were obtained from two experimental replicates.

**Supplemental Figure S11.**
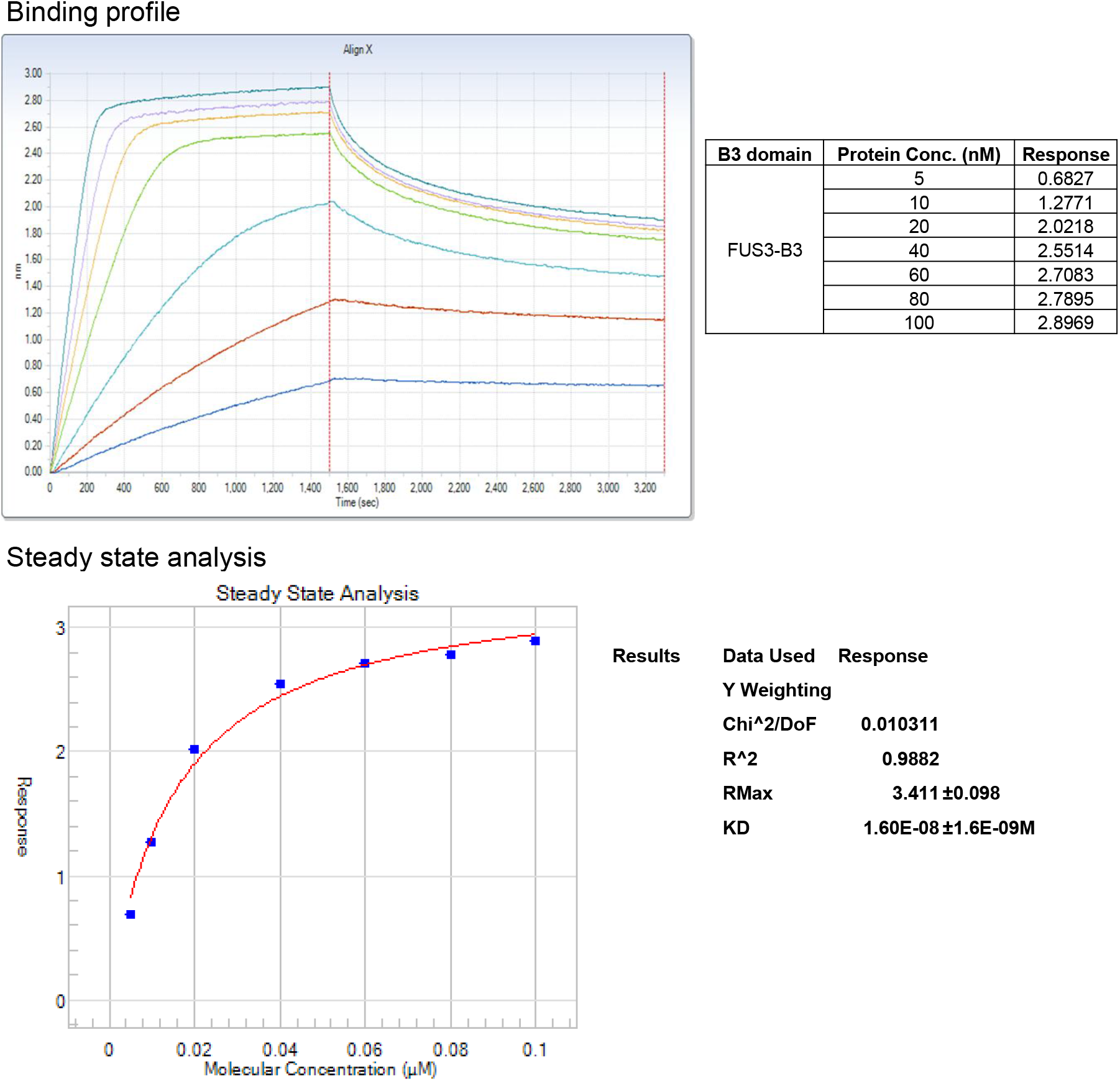
Quantification of FUS3-B3 domain DNA binding activity by Octet Qke. B3-DNA-binding was measured in a label-free in vitro kinetics assay at pH 7.0 using Octet Qke system (Pall ForteBio). Experimental conditions and data analysis were described in the Supplemental Figure S10 and materials and methods.

**Supplemental Figure S12.**
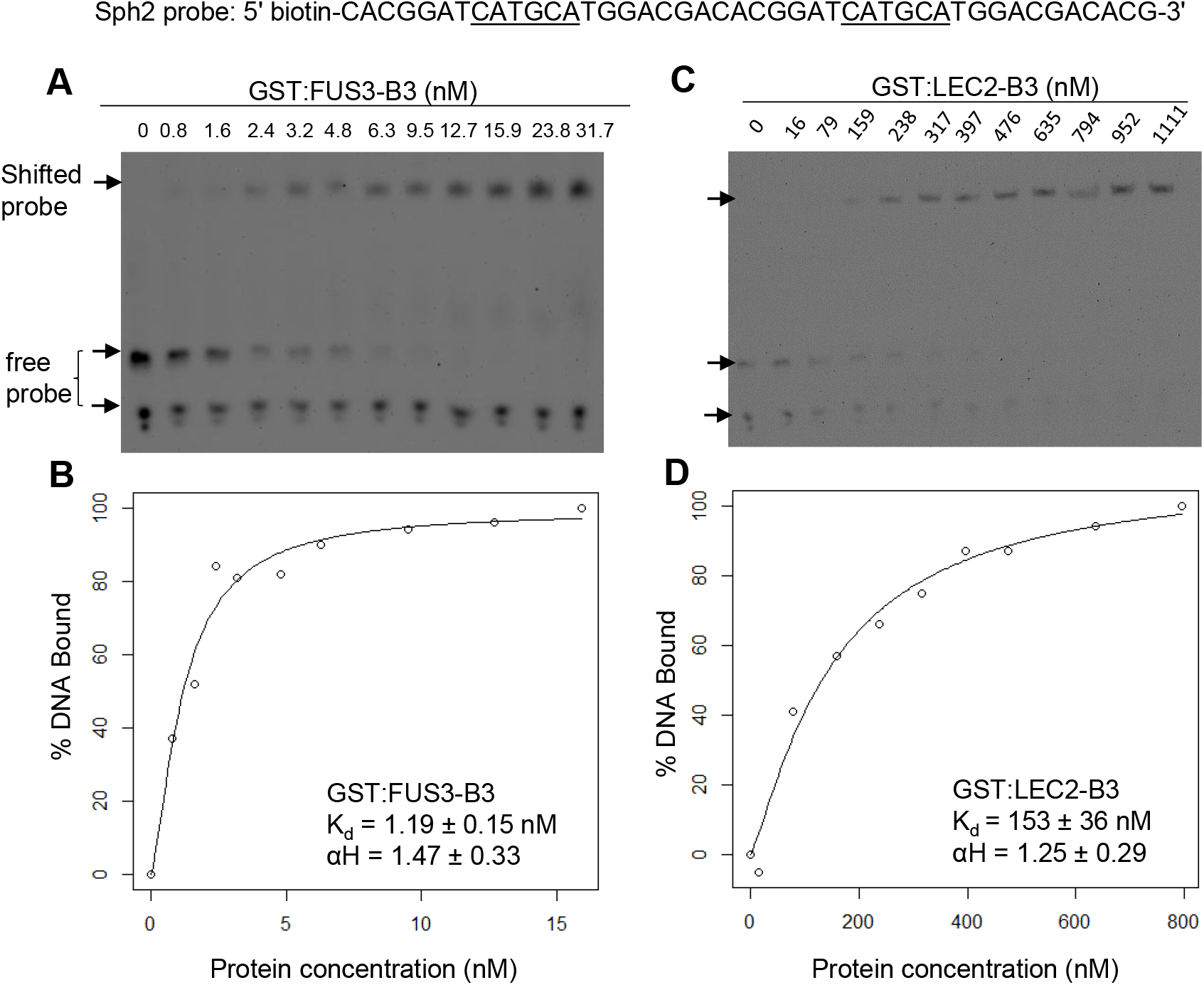
Analysis of B3 domain DNA binding activity by gel mobility shift assay. GST:B3 binding to the Sph2 DNA probe was determined by gel mobility shift assays varying protein concentration. The probe sequence is shown at the top of the figure. Chemiluminescence images in (**A**) and (**C**) were captured by a CCD camera. The fraction of the total probe shifted was quantified by densitometry using image J. Binding curves GST:FUS3-B3 (**B**) and GST:LEC2-B3 (**D**) and associated kinetic parameters were determined by nonlinear least squares fitting the equation Y = B_max_ * X^αH^/ (K_d_^αH^ + X^αH^), where Y is the percentage of total probe shifted, X is the protein concentration in nM units, αH is the Hill constant, and K_d_ is the dissociation constant (in nM units).

**Supplemental Figure S13.**
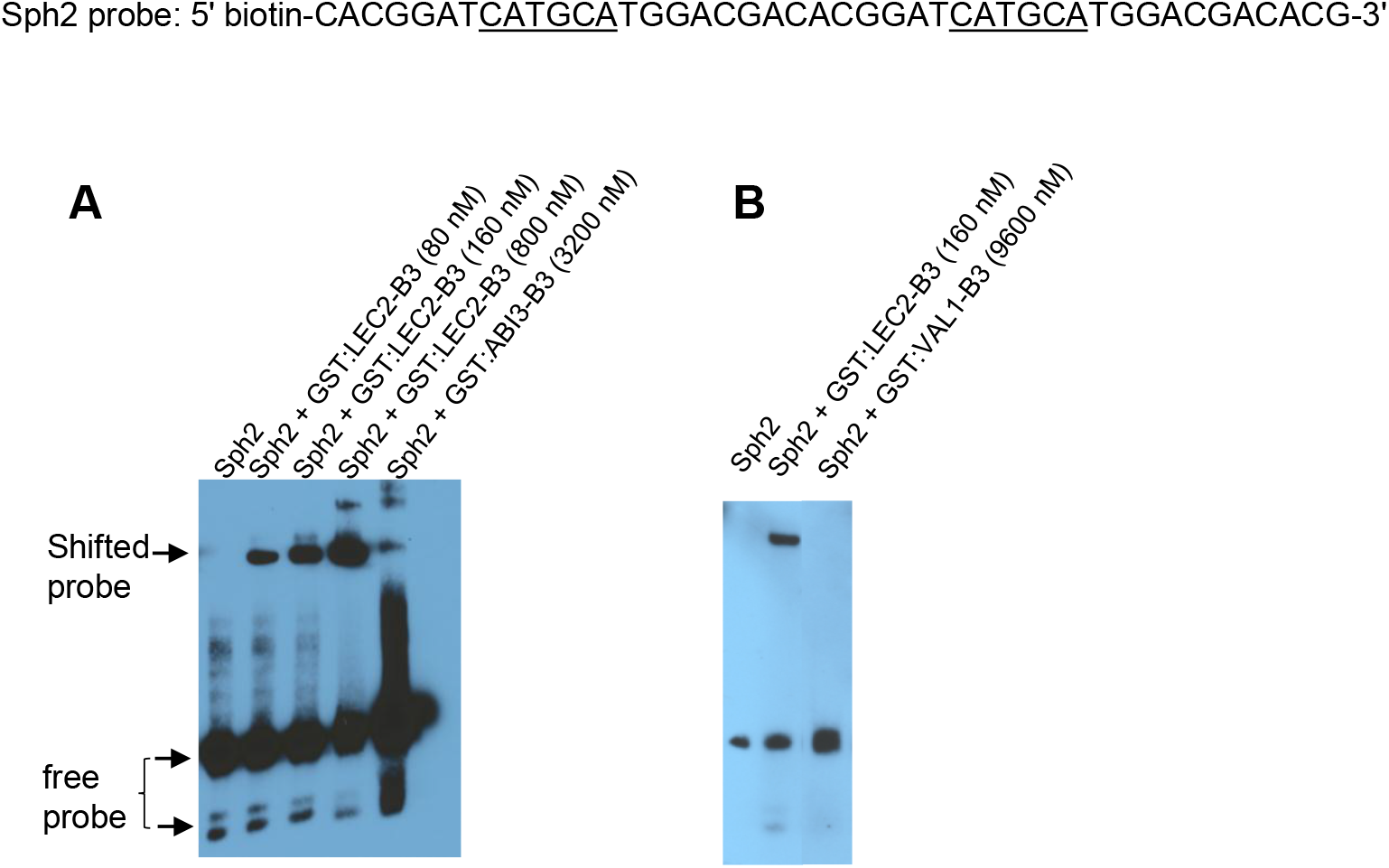
Analysis of ABI3 and VAL1 B3 domain DNA binding activity by gel mobility shift assay. (**A**) ABI3 B3 domain was tested in the same condition as LEC2 B3 domain (**B**) VAL1 B3 domain was tested in the same condition as LEC2 B3 domain. Experimental conditions were described in the Supplemental Figure S12 and materials and methods.

**Supplemental Figure S14.**
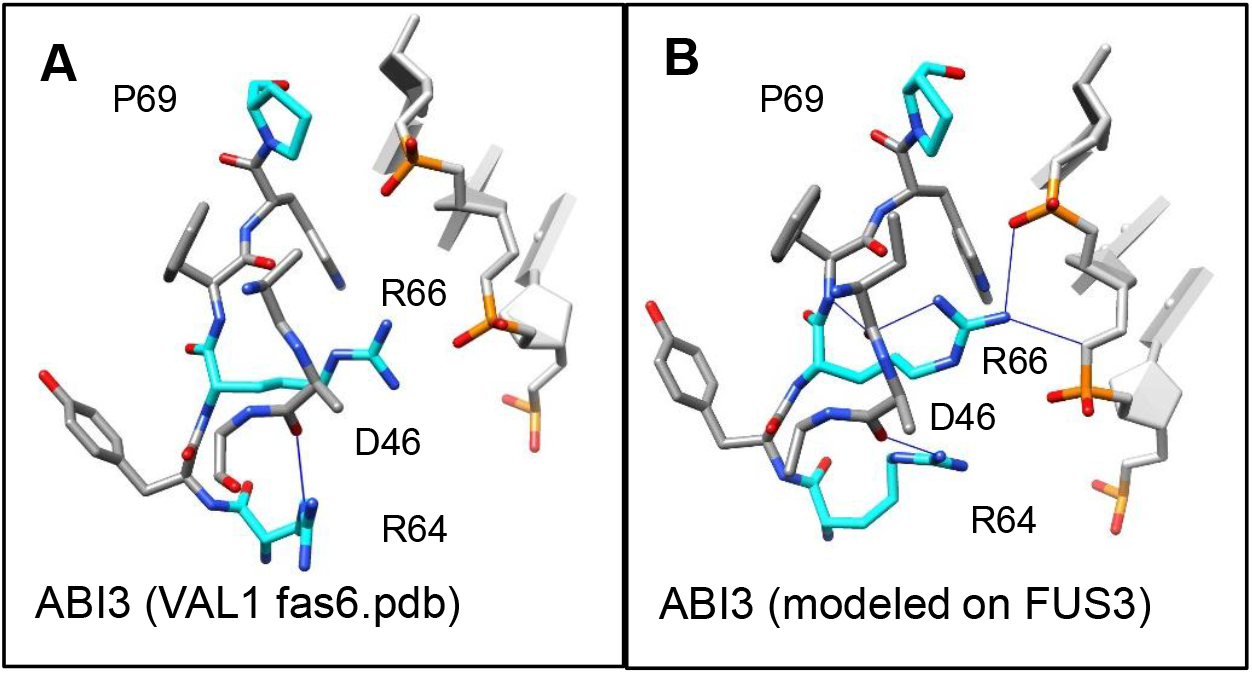
H-bonding of R64 with the backbone carbonyl of D46 in independent ABI3 models. (**A, B**) ABI3 models based on the Sasnauskas et al. VAL1 structure (6FAS.pdb) (8) and FUS3 (6J9B.pdb) (9). H-bonds (blue lines) were identified using the findHBond tool in Chimera (18) with relaxed constraints.

**Supplemental Table S1.**
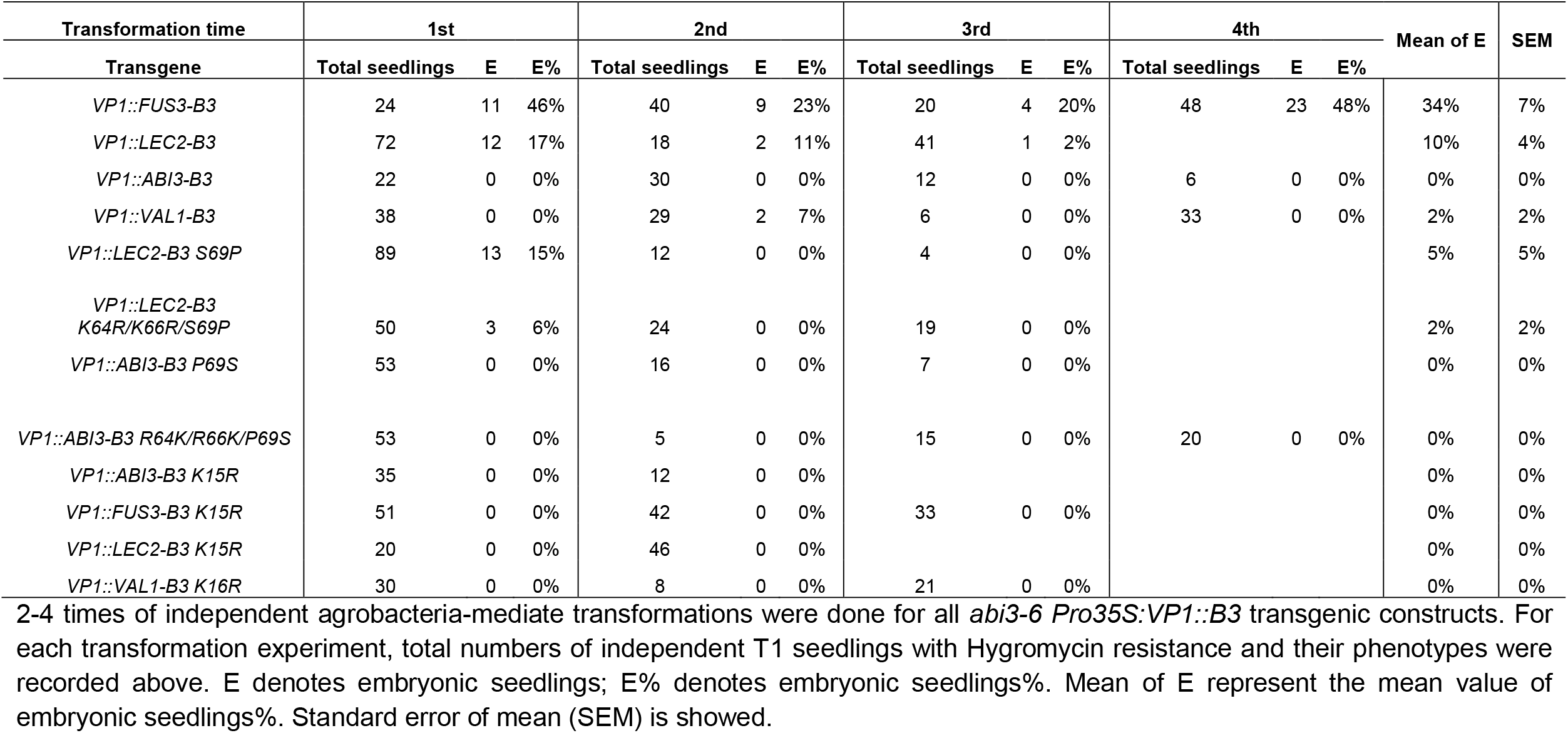
Phenotypes of *abi3-6 Pro35S:VP1::B3* T1 Transgenic Plants

| Transformation time | 1st |  |  | 2nd |  |  | 3rd |  |  | 4th |  |  | Mean of E | SEM |
| --- | --- | --- | --- | --- | --- | --- | --- | --- | --- | --- | --- | --- | --- | --- |
| Transgene | Total seedlings | E | E% | Total seedlings | E | E% | Total seedlings | E | E% | Total seedlings | E | E% |  |  |
| *VP1::FUS3-B3* | 24 | 11 | 46% | 40 | 9 | 23% | 20 | 4 | 20% | 48 | 23 | 48% | 34% | 7% |
| *VP1::LEC2-B3* | 72 | 12 | 17% | 18 | 2 | 11% | 41 | 1 | 2% |  |  |  | 10% | 4% |
| *VP1::ABI3-B3* | 22 | 0 | 0% | 30 | 0 | 0% | 12 | 0 | 0% | 6 | 0 | 0% | 0% | 0% |
| *VP1::VAL1-B3* | 38 | 0 | 0% | 29 | 2 | 7% | 6 | 0 | 0% | 33 | 0 | 0% | 2% | 2% |
| *VP1::LEC2-B3 S69P* | 89 | 13 | 15% | 12 | 0 | 0% | 4 | 0 | 0% |  |  |  | 5% | 5% |
| *VP1::LEC2-B3 K64R/K66R/S69P* | 50 | 3 | 6% | 24 | 0 | 0% | 19 | 0 | 0% |  |  |  | 2% | 2% |
| *VP1::ABI3-B3 P69S* | 53 | 0 | 0% | 16 | 0 | 0% | 7 | 0 | 0% |  |  |  | 0% | 0% |
| *VP1::ABI3-B3 R64K/R66K/P69S* | 53 | 0 | 0% | 5 | 0 | 0% | 15 | 0 | 0% | 20 | 0 | 0% | 0% | 0% |
| *VP1::ABI3-B3 K15R* | 35 | 0 | 0% | 12 | 0 | 0% |  |  |  |  |  |  | 0% | 0% |
| *VP1::FUS3-B3 K15R* | 51 | 0 | 0% | 42 | 0 | 0% | 33 | 0 | 0% |  |  |  | 0% | 0% |
| *VP1::LEC2-B3 K15R* | 20 | 0 | 0% | 46 | 0 | 0% |  |  |  |  |  |  | 0% | 0% |
| *VP1::VAL1-B3 K16R* | 30 | 0 | 0% | 8 | 0 | 0% | 21 | 0 | 0% |  |  |  | 0% | 0% |

2-4 times of independent agrobacteria-mediate transformations were done for all *abi3-6 Pro35S:VP1::B3* transgenic constructs. For each transformation experiment, total numbers of independent T1 seedlings with Hygromycin resistance and their phenotypes were recorded above. E denotes embryonic seedlings; E% denotes embryonic seedlings%. Mean of E represent the mean value of embryonic seedlings%. Standard error of mean (SEM) is showed.

**Supplemental Table S2.**
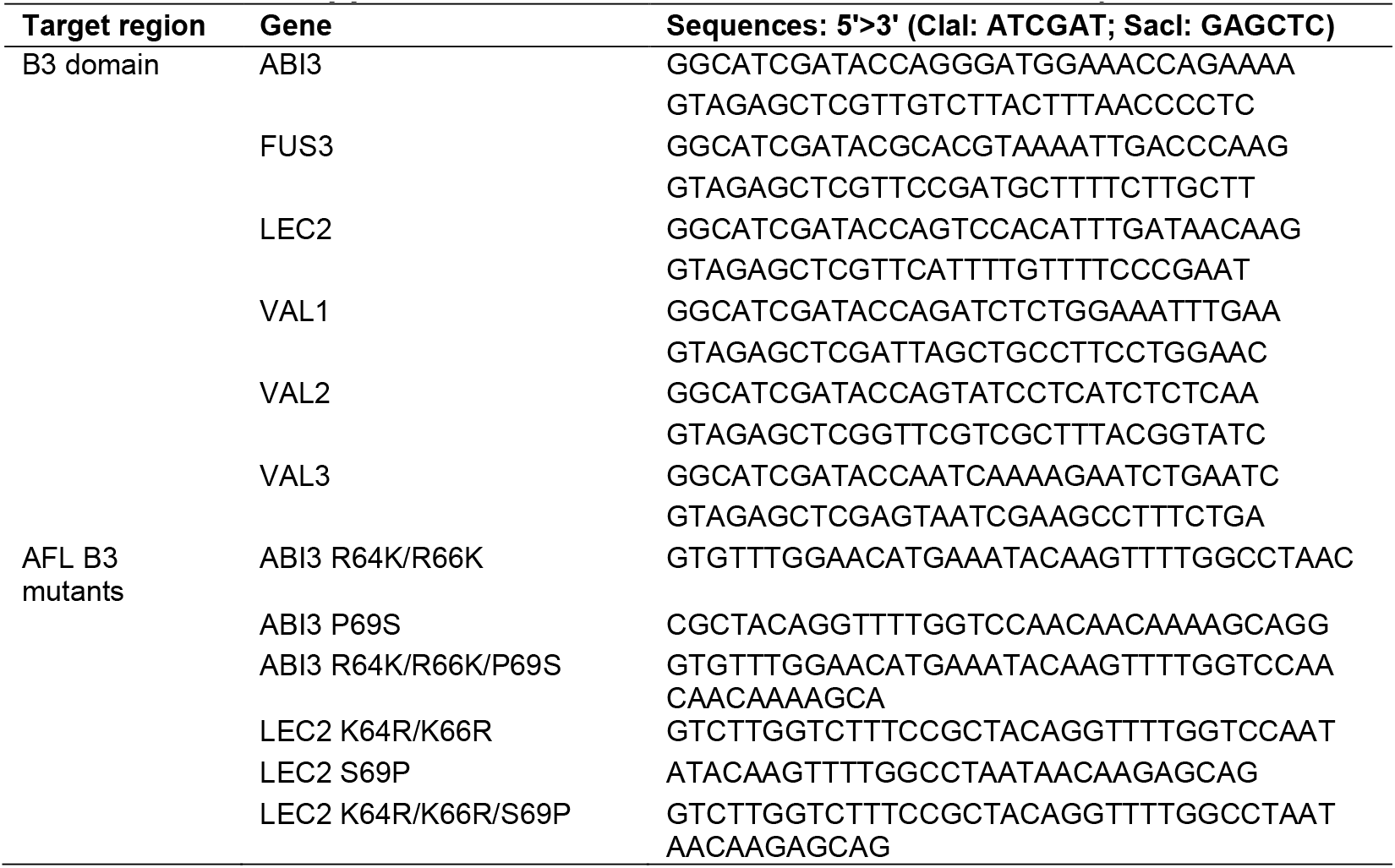
Primers used in Y1H study

**Supplemental Table S3.**
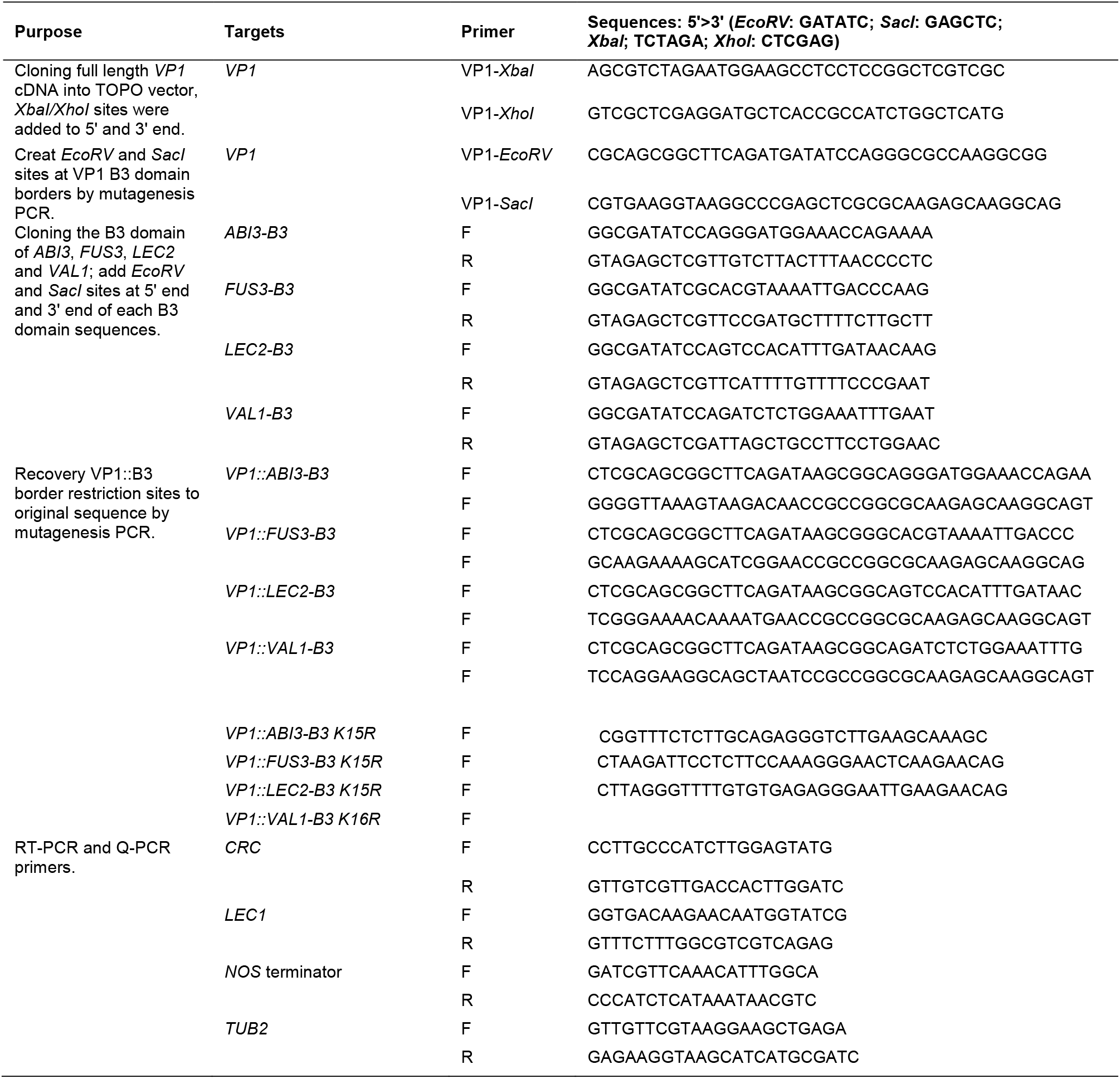
Primers used in transgene construction and transgenic line analysis

**Supplemental Table S4.**
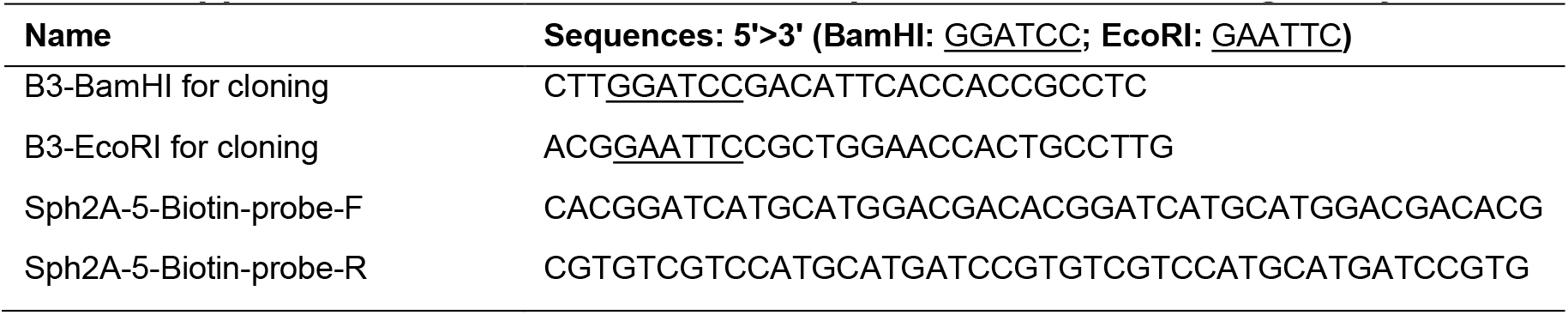
Primers and probes used in binding assays

**Supplemental Table S5.**
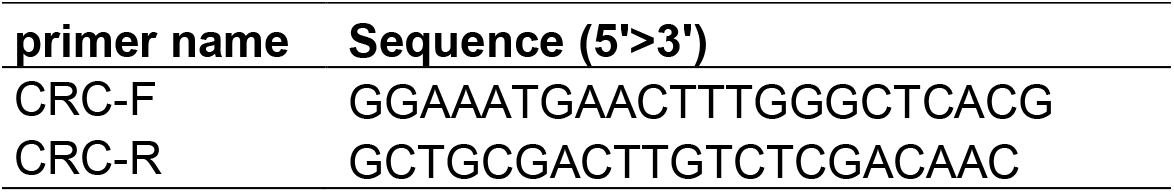
Primers for ChIP-qPCR

